# Phenome-wide analysis identifies parent-of-origin effects on the human methylome associated with changes in the rate of aging

**DOI:** 10.1101/2023.01.18.524653

**Authors:** Chenhao Gao, Carmen Amador, Rosie M. Walker, Archie Campbell, Rebecca A Madden, Mark J. Adams, Xiaomeng Bai, Ying Liu, Miaoxin Li, Caroline Hayward, David J. Porteous, Xueyi Shen, Kathryn L. Evans, Chris S. Haley, Andrew M. McIntosh, Pau Navarro, Yanni Zeng

## Abstract

Variation in the rate at which humans age may be rooted in early life events acting through genomic regions that are influenced by such events and subsequently are related to health phenotypes in later life. The parent-of-origin-effect (POE)-regulated methylome includes regions either enriched for genetically controlled imprinting effects (the typical type of POE) or atypical POE introduced by environmental effects associated with parents. This part of the methylome is heavily influenced by early life events, making it a potential route connecting early environmental exposures, the epigenome and the rate of aging. Here, we aim to test the association of POE-influenced methylation of CpG dinucleotides (POE-CpG sites) with early and later environmental exposures and subsequently with health-related phenotypes and adult aging phenotypes. We do this by performing phenome-wide association analyses of the POE-influenced methylome using a large family-based population cohort (GS:SFHS, N_discovery_=5,087, N_replication_=4,450). At the single CpG level, 92 associations of POE-CpGs with phenotypic variation were identified and replicated. Most of the associations were contributed by POE-CpGs belonging to the atypical class and the most strongly enriched associations were with aging (DNAmTL acceleration), intelligence and parental (maternal) smoking exposure phenotypes. We further found that a proportion of the atypical-POE-CpGs formed co-methylation networks (modules) which are associated with these phenotypes, with one of the aging-associated modules displaying increased internal module connectivity (strength of methylation correlation across constituent CpGs) with age. Atypical POE-CpGs also displayed high levels of methylation heterogeneity and epigenetic drift (i.e. information loss with age) and a strong correlation with CpGs contained within epigenetic clocks. These results identified associations between the atypical-POE-influenced methylome and aging and provided new evidence for the “early development of origin” hypothesis for aging in humans.

## Introduction

Aging is a multi-system process manifesting as a progressive decline of physiological integrity, impaired functions and increased risk of adult onset diseases and death(1). Although everyone ages chronologically, the actual biological state, namely biological age, varies even among individuals of the same chronological age(2, 3). The increased or delayed biological aging after accounting for chronological age has been defined as “age acceleration”, which can be estimated by biomarkers such as DNA methylation(4–7). Identification of risk factors and biomarkers is crucial for the understanding of aging(2). Genetic studies have reported large numbers of genomic loci associated with biological aging(8). Biological aging explained by DNA sequences, however, only accounts for influences from predisposing and unchangeable risk factors. Environment-involved effects such as epigenetic changes in response to life events, on the other hand, are flexible and reversible, representing a different collection of factors which could potentially better fit the dynamic nature of aging process across the lifespan.

Among all environmental factors, early and developmental exposures are of particular interest. In 1994, Barker proposed a hypothesis that late-onset disease can be profoundly influenced by early life experiences(9). Since then, a number of studies have provided “early development of origin” evidence for adult-onset diseases such as schizophrenia and dementia(10, 11). Aging, which is the biggest risk factor for many late onset diseases, has been found to be associated with adulthood environmental factors such as smoke and sun exposures(12, 13). When it comes to early effects, a few studies reported association between early exposures and age acceleration in newborns and children, but not in adults (not tested)(13, 14). The association between early events/exposures and adulthood aging, and the molecular paths mediating any such associations, have been largely unexplored.

Parent-of-origin effects (POEs) are found in a subset of genomic regions that are highly sensitive to early-life events and associated with health outcomes at both early and late life stages(15–17). At the level of DNA methylation, typical POE-influenced methylation sites manifest as imbalanced methylation similarity between nuclear family members of the same genetic distance (mother-offspring, father-offspring, sibling-sibling pairs) and differentiated allelic effect on methylation levels depending on the parent of origin of the regulatory SNPs (POE-mQTLs)(16, 18). This type of POE is mainly introduced by genomic imprinting, which is established at early developmental stages and needs to be well maintained/regulated throughout the life(16). The epigenomic features influenced by typical POEs has been found to be sensitive to prenatal and postnatal environmental stimuli, such as maternal nutrition during pregnancy and stress accompanied with assisted reproductive technologies(15, 19–22). In addition to the methylation sites influenced by the typical POE, a different set of methylation sites also display imbalanced methylation similarity between nuclear family members of the same genetic distance, but without regulatory POE-mQTLs being detected, and are not enriched in known imprinted regions. These sites should, therefore, be regarded as “atypical POE-CpGs”(18). Since dominance effects have been ruled out for the majority of these atypical POE-CpGs(18), the potential explanations for the atypical POE pattern are either small POE-mQTL (imprinting) effect not yet detected due to the lack of power, or early familial environmental effects introduced by the parents(18). In any case, both typical and atypical POE-CpGs represented classes of CpGs where methylation levels are heavily influenced by early life events. Once involved in physiological functions in later life, they can be pivotal to the interplay between early-life experiences, epigenome, and adulthood health(15, 23).

The early-life-event-sensitive nature of the POE-regulated methylome renders it a plausible mechanism for the “early development of origin” hypothesis of adult aging. The link between POE and aging has been suggested by a few animal studies including one showing that the knockout of imprinted gene *RasGrf1* promoted longevity(24); and further two showing that early-life adversity caused imprinting deregulation of gene *Cdkn1c*, resulting in interrupted expression which influences aging-associated obesity(25, 26). Human studies on association between POE and aging, however, are very limited. POE studies in humans mostly targeted rare developmental diseases (mainly imprinted disorders) caused by genetic mutation, other studies mainly examined genetic effects that influence complex traits in a parent-of-origin way(27–29). These included studies focused on late-onset disease such as Alzheimer’s disease(30), but few have studied aging phenotypes (such as age acceleration) themselves. Moreover, even to examine aging phenotypes in future studies, the genotype-based strategies which the majority of existing human studies rely on do not account for the environment-sensitive and dynamic features of POE-influenced genomic regions, which may lead to underestimation of the effects from these regions. Methylation studies, on the contrary, capture effects from both genetic background and environmental exposures, offering unique advantages in this context. To date, only one human study reported the association between methylation levels of POE-influenced genes and the change of brain structures overtime, but with a relatively small sample size (N=485) and only investigated a small proportion of POE-influenced genes (13 imprinted locations) (31). Therefore, a well-powered systematic examination of associations between POE-regulated methylome and adult aging is warranted.

In this study, we aimed to investigate POE-influenced methylome to collect evidence for the “early development of origin” hypothesis for aging in humans (Figure 1). At both single CpG and co-methylation network levels, the associations between 943 POE-CpGs (N_typical_=560, N_atypical_=383) and 142 phenotypes were tested and replicated using two subsets of GS:SFHS (Generation Scotland: Scottish Family Health Study. N_discovery_=5,081, N_replication_=4,445), a large family-based population cohort with genome-wide DNA methylation data (N_sites_=734,436), records of early to late life exposures and extensive health-related phenotypes available for participants(32, 33). The phenotypes included four aging measurements: two epigenetic-based acceleration variables - DNAmTL acceleration and PhenoAge acceleration, and parental lifespans. As aging is the underlying cause of many adulthood illnesses, we expected widespread associations between aging-associated POE-CpGs with health-related phenotypes, therefore a phenome-wide scan was applied instead of only testing for a few aging phenotypes. Our primary results revealed strongly enriched associations of atypical type POE-CpGs with early and late life exposures and with aging-related phenotypes at both single CpG and co-methylation network level. An aging-associated atypical POE co-methylation module whose internal connectivity increased with age was further identified. These findings motivated two additional aging-focused analyses, which revealed high levels of methylation heterogeneity and epigenetic drift in atypical POE-CpGs and intrinsic connections between atypical POE CpGs and clock CpGs (Figure 1).

**Figure 1.**
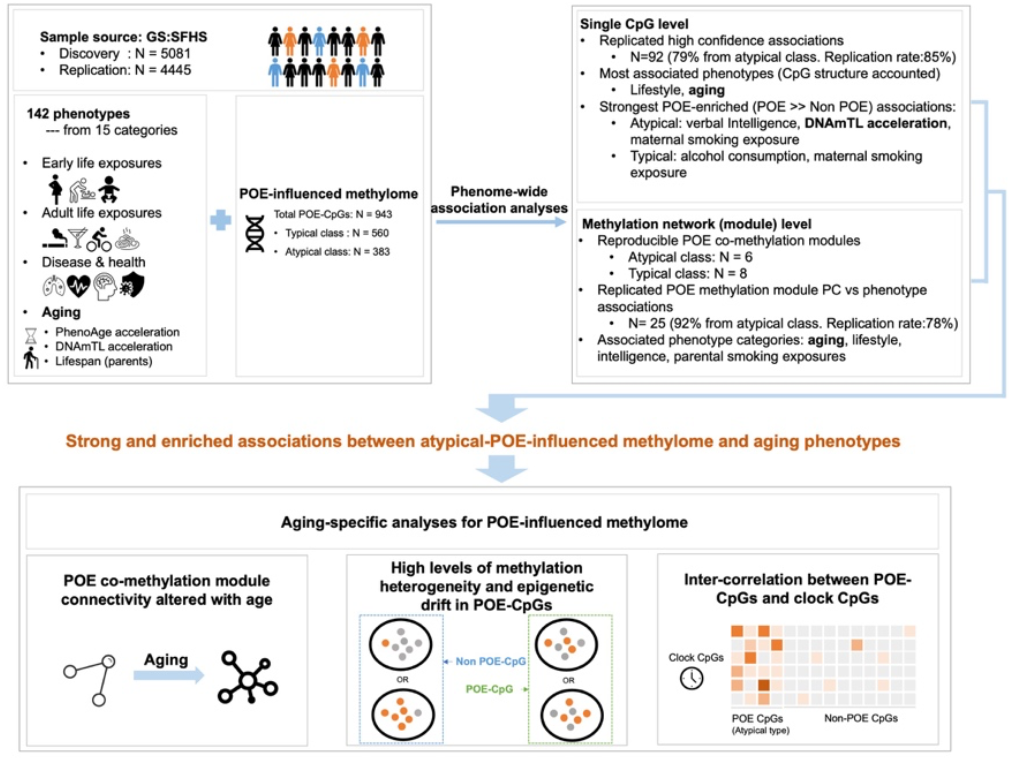
An overview of the study design.

## Methods

### Ethics

The ethical approval of GS:SFHS studies was obtained from the Tayside Research Ethics Committee (reference 05/S1401/89). Before any data or samples were collected, participants all gave written consent after having an opportunity to discuss the research.

### Population samples

GS:SFHS is a family-based population cohort with extensive health-related phenotypes, records of environmental exposures, genome-wide genotypes collected for 19,994 Scottish participants(32, 34). Genome-wide DNA methylation data (whole blood) was also available for 9,526 participants(33). The methylation data was produced and processed independently in two batches, for 5,081 participants in 2016-2017 (batch 1) and 4,445 participants in 2019 (batch 2). All participants in batch 2 were genetically unrelated (relatedness < 0.05) to each other and to the participants in batch 1. We used batch 1 as discovery dataset and batch 2 as replication dataset in downstream analyses.

### DNA methylation data

The discovery and replication datasets were generated, processed and quality controlled in a similar way (35) based on a pipeline proposed previously (18, 36). In brief, methylation signals for 866,836 sites were measured using the Illumina Infinium MethylationEPIC array (http://support.illumina.com) for whole blood sample of each participant. The “estimateCellCounts” function in R package minfi was used to estimate the proportion of major blood cell types: B-lymphocytes, natural killer cells, monocytes, granulocytes, CD4+ T-lymphocytes and CD8+ T-lymphocytes (37). The R packages shinyMethyl and meffil were used for quality control (38, 39). The performance of control probes, signal intensity, and consistency between registered and predicted sex were used to identify outlier samples and probes. In addition, samples were removed if more than 0.5% of measured sites had a detection p-value > 0.01. Probes were removed if more than 1% of samples were missing or had a bead count ≤ 3, or if they had cross-hybridization or overlapped with any common SNP (MAF≥0.01) in the European population (40). After quality control, normalization was performed using the “ssNoob” method in R package minfi (41). As described before(18), normalized M values were adjusted, using a linear mixed model, for technical variables including sentrix variables (id and position), processing batches, clinics, appointment variables for the blood extraction(date, weekday, and year), and 20 principal components calculated from the control probes (36). Resultant residuals were available for 734,436 methylation sites which were used in downstream analyses.

### Collection of POE-influenced methylation sites

The list of 984 POE-CpGs (N_atypical_=398, N_typical_=586) was extracted from a previous study(18). After quality control for DNA methylation data (see above), 943 POE-CpGs (N_atypical_=383, N_typical_=560) were used in downstream analyses.

### Phenotype data

Phenotypes in GS:SFHS consisted of 142 variables in 15 categories (Table s1). Among them, birth and maternity variables were obtained through data linkage with historic Scottish birth cohorts for a subset of GS:SFHS participants(42). The aging category comprised four variables, including mother’s/father’s lifespan and two epigenetic-based measurements for biological aging (PhenoAge acceleration and DNAmTL acceleration) (6, 7). The two acceleration measurements were calculated as the residuals from regressions of PhenoAge, an epigenetic clock designed to predict healthspan (phenotypic age) (6), and DNAmTL, an epigenetic clock designed to predict telomere length(7), on age and age^2^. Positive PhenoAge acceleration corresponds to excessive biological aging among individuals of the same chronological age, whereas positive DNAmTL acceleration corresponds to the additional (longer) telomere length after accounting for chronological age. The phenotypic correlation between the four aging measures is shown in Figure s1. Quantitative traits with a skewed distribution were log transformed with base 10. Measurements that fall outside of four standard deviations from the mean were identified as outliers and thus removed. More details of the phenotypes are provided in Table s1

### Phenome-wide association analyses for POE-influenced methylation sites

Phenome-wide association analyses for individual POE-CpG sites were performed using MLM-based Omic association (MOA) models:

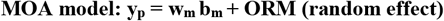

As proposed by the Omic-data-based Complex trait Analysis software (OSCA)(43), MOA is a linear mixed modelling method that allows adjusting for the global correlation between probes which is likely introduced by unobserved confounders. This was realized by fitting an Omic-Relationship-Matrix (ORM) as a random effect jointly with the target CpG variable as a fixed effect in linear mixed models(43). The ORM represented the epigenetic relationships between samples, created by the “--make-orm” function in OSCA using genome-wide probes (N=734,436. M values were pre-adjusted for cell proportion, appointment variables, age, age^2^, sex, and smoking variables (smoking status and pack years). Age and age^2^ were not preadjusted if PhenoAge acceleration or DNAmTL acceleration was the target phenotype; smoking variables were not pre-adjusted if smoking status was the target phenotype). **y_p_** was the target phenotype pre-adjusted for two random effects represented by the genomic relationship matrix (G) and the kinship relationship matrix (K) (accounting for the genetic structure in GS:SFHS), and clinic effect (as fixed effect), using genome-based restricted maximum likelihood (GREML) in GCTA(44). **w_m_** was the methylation level of the target CpG site after pre-adjusting for blood cell proportion, appointment variables, age, age^2^, sex and smoking variables as fixed effects (age and age^2^ were not pre-adjusted when PhenoAge acceleration or DNAmTL acceleration was the target phenotype; smoking variables were not pre-adjusted when smoking status was the target phenotype). **b_m_** was the target effect to be estimated.

The MOA approach was applied to each of the POE-CpG and phenotype pairs. Since the pre-adjustment did not converge for 9 out of the 943 POE-CpGs, we only included the remaining 934 POE-CpGs in this analysis. The false discovery rate (FDR) method was used to correct for multiple testing in both discovery and replication stages (N_tests_discovery_=934*142=132628, N_tests_in_replication_=N_significant_pairs_in_discovery_). For replicated results, we additionally used a classical linear regression model which avoids pre-adjustment of methylation-related variables to further validate the robustness of the MOA results:

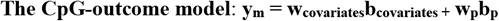

In contrast to the MOA models, the CpG-outcome model is a linear fixed-effect regression model that takes methylation levels of the target CpG sites as the dependent variable and the target phenotype values as the independent variable, with methylation-related biological covariates being jointly fitted in the model to avoid having to pre-adjust for those covariates. **y_m_** was the methylation level of target CpG sites after pre-adjusting for the G and K components as random effects (to account for genetic structure) and clinic effect as a fixed effect using GREML(44). **w_covariates_** was a matrix for covariates including blood cell proportion, appointment variables, age, age^2^, sex and smoking variables (age and age^2^ were not fitted when PhenoAge acceleration or DNAmTL acceleration was the target phenotype; smoking variables were not fitted when smoking status was the target phenotype). b_covariates_ was the effects from covariates. **w_p_** was the target phenotype and **b_p_** was the target effect to be estimated. We only considered the results that were statistically significant and replicated in both the MOA model and the CpG outcome model as high confidence results.

### Comparison of phenotypic associations with POE vs Non-POE methylome

This analysis was to test whether for a given phenotype, its association with POE-CpGs was significantly stronger than its associations with the rest of methylome. In brief, methylome-wide association studies (MWASs, N_CpG_=734,438) were performed using the same MOA approach for phenotypes associated with at least one POE-CpG. The Wilcoxon Rank Sum Test was then applied to each phenotype to test whether the P values of POE-CpG-specific methylation-phenotype associations were ranked significantly differently from the P values of the rest of methylome. The Bonferroni method was applied to adjust for multiple testing correction.

### Identification of modules of co-methylated CpGs in the POE-influenced methylome

Weighted gene correlation network analysis(WGCNA) was applied to identify modules of co-methylated POE-CpGs (45). Before constructing modules, methylation levels of POE-CpGs were pre-corrected by cell proportions, appointment variables, age, age^2^, sex and smoking variables.

Given the differentiated features of typical and atypical POE-CpGs, co-methylation modules were constructed for typical-type (N=560) and atypical-type (N=383) POE-CpGs separately, and for discovery (only unrelated samples (relatedness < 0.05) were used in network construction, N=2,583) and replication datasets separately. The “soft thresholding power” parameter was optimized to allow identification of both tightly-connected CpG clusters such as those in *cis* (for example, CpGs from the same island) and modestly-connected CpG clusters such as those in *trans* (for example CpGs in different chromosomes). In detail, a recursive process was applied as following: 1) all typical/atypical POE-CpGs were used to fit the “PickSoftThreshold” function and construct networks. In this step the picked threshold was high and only tightly connected CpGs could be assigned to modules. 2) For each module identified by step 1, only one hub CpG displaying the highest correlation with other CpGs was retained in every 10 kb window. 3) Step 1 and 2 were repeated until no more typical/atypical POE-CpGs were removed. 4)The retained set of typical/atypical POE-CpGs was used to re-fit the PickSoftThreshold function. At this stage, the optimised soft thresholding power could be estimated. 5) We used this optimised parameter (equal to three) to construct full networks using all typical/atypical POE-CpGs. The smallest number of CpGs for a module was set to 8. Other parameters were set to the default ones.

### Matching POE co-methylation modules across discovery and replication datasets

Since POE co-methylation modules were identified independently in discovery and replication datasets, we matched modules in the two datasets using following steps: 1) for any two modules, one from the discovery dataset and one from the replication dataset, the overlap rate was calculated as the number of CpGs in the intersection divided by the number of CpGs in the union. 2) All discovery-replication module pairs were ranked by overlap rate in descending order. Starting from the top pair, if the overlap rate was higher than 60%, modules across datasets were successfully matched. 3) For modules identified in the replication dataset but not matched with any module in the discovery dataset yet, we calculated the secondary overlap rate with each discovery module, defined as the number of CpGs in the interaction divided by the number of CpGs in the replication module. The replication module could be matched to the discovery module (that is, to allow more than one replication modules to be matched to one discovery module) if the secondary overlap rate is higher than 90%. 4) Matched modules were labeled as “consistent modules”, with shared CpGs labeled as constituent CpGs which were used in downstream analyses.

### Phenome-wide association analyses for POE co-methylation modules

#### Identification of principal components for POE co-methylation modules

To quantify POE co-methylation modules, we performed principal component analysis (PCA) for methylation levels of constituent CpGs for each “consistent module” using unrelated samples (relatedness < 0.05) from discovery dataset (N=2,583). The estimated formula was then projected to the entire cohort to calculate module PCs for all discovery and replication samples. This was done by the R package “psych” (https://CRAN.R-project.org/package=psych). In downstream analyses, we only used module PCs that explained >5% of the methylation variation of the corresponding module. Similar to the single-CpG based analyses, analyses for phenotypes such as the two age acceleration phenotypes and smoking status required a modified list of covariates. We therefore prepared three sets of PCs by using methylation levels pre-corrected for different sets of covariates:

PC set 1: pre-corrected for cell proportions, appointment variables, smoking variables, age, age^2^, sex.
PC set 2: same as PC set 1 except without pre-correcting for smoking variables.
PC set 3: same as PC set 1 except without pre-correcting for age and age^2^.

#### Phenome-wide association tests for POE co-methylation module PCs

Linear regression was used to regress module PCs on the phenotype:

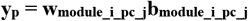

Similar to single-CpG-based tests, y_p_ represented target phenotypes pre-adjusted for G and K components as random effects and clinic as fixed effect. **w_module_i_pc_j_** was the the top ith PC in module j. **b_module_i_pc_j_** was the tested effect from the ith PC of module j. Since methylation-related covariates have been pre-adjusted when generating the PCs (described above), we did not re-adjust for covariates at this step. The module PC set 1 (described above) was used for most association tests, except for the tests targeting smoking status (the module PC set 2 was used), and the tests targeting age acceleration phenotypes (the module PC set 3 was used).

### Analyses of the dynamics of internal methylation connectivity for aging-associated POE co-methylation modules across age-stratified groups

We stratified samples into six subgroups according to their chronological age (Table s2). For each aging-associated POE co-methylation module, connectivity among constituent CpGs was measured using pairwise Pearson correlation of methylation levels pre-adjusted for cell proportion, sex, appointment variables and smoking variables. The overall distribution of connectivity across age groups was tested by the Kruskal–Wallis method using R function “kruskal.test”, the connectivity difference between any two age groups was tested by the Wilcoxon Rank Sum Test using R function wilcox.test, and the variance of connectivity across age groups were tested by the Levene’s test using function “levenetest” in R package “car”(46). Based on age-dependent connectivity trajectories, sub clusters within the module of interest were identified using hierarchical clustering. Cytoscape was used to calculate the node centrality and visualize the results(47).

### OMIC- and Summary-data-based Mendelian Randomization (SMR) analysis

SMR was applied to identify pleiotropic associations between methylation levels of target CpG and mRNA expression levels of nearby genes(48). Brain *cis*-mQTL summary statistics were from Qi et al., 2018(49), brain *cis*-eQTL summary statistics were from Qi et al., 2022 (unpublished, the data (BrainMeta v2) was accessed through the software SMR(48)). In this analysis, methylation was treated as the exposure and mRNA expression was treated as the outcome. The Bonferroni method was applied to correct for multiple testing in SMR analyses. HEIDI test was applied to distinguish pleiotropy from linkage, with a P_HELDI_ > 0.05 (unadjusted) indicating that the association was not due to linkage(48).

### Permutation tests for the connectivity between clock CpGs and POE-CpGs

The lists of CpGs used in the construction of two first-generation epigenetic clocks, Hannum and Horvath clocks, and two second-generation epigenetic clocks, PhenoAge and DNAmTL, were downloaded from the original publications, respectively(4–7). Circular permutation over the methylome was used to generate 10,000 random CpG sets of the same size as the typical/atypical POE-CpGs, keeping the overall correlation structure of the true POE-CpG set in the generated random sets(50). For each clock, the average connectivity between clock CpGs and POE-CpGs (the true set and the permuted sets) was calculated as the mean of absolute values of pairwise methylation correlation (Pearson method). Permutation P values were calculated by ranking the average connectivity of permuted sets in descending order and determining the position of the true average connectivity in the ranked list.

### Calculation of methylation Shannon entropy

For DNA methylation, the Shannon entropy measures the level of methylation uncertainty (methylation heterogeneity)(5, 51, 52). The following formula was used to calculate the Shannon entropy for a given CpG in a given sample(52):

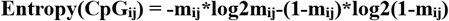

Where **m_i_** is the Beta value of a given CpG i for a given sample j.

### Annotation and visualization

Functional annotations for CpGs and genes were performed using ANNOVAR(53). The R packages ggplot2(54), ggpubr(55), ComplexHeatmap(56) and visNetwork(57) were used in the visualization of the presented results.

## Results

### Phenome-wide association analyses identified strong and enriched associations of atypical POE-CpGs with aging, intelligence and early/late environmental exposures

The MOA method was applied to test the association between each POE-CpG and each phenotype (N_CpG_=934, N_phenotype_=142). At the discovery stage (N_sample_=5,081), a total of 115 POE-CpG-phenotype pairs exceeded the phenome-wide significant threshold (FDR < 0.05). At the replication stage (N_sample_=4,445), 85.2% (N=98) of the POE-CpG-phenotype associations were statistically replicated (FDR < 0.05) (Table s3). The CpG-outcome model further validated the robustness of 94% (N=92) of the replicated associations reported by the MOA method (Table s3), we considered this set as “high confidence associations” (Table s4).

The 92 high confidence associations involved 38 POE-CpGs and 24 phenotypes, revealing widespread associations of POE-CpGs with multiple phenotype-categories (Figure 2). The majority of the associations (79.3%) was contributed by atypical POE-CpGs (Table s4), despite the fact that the atypical group only accounted for 40.6% of the total POE-CpGs. The phenotypic categories contributed the largest number of associations were parental smoking exposure, lifestyle, intelligence and aging (Figure 3a). Conditional analyses suggested that the associations with parental smoking were mainly driven by maternal smoking (Table s5). After accounting for the baseline number of POE-CpGs in the atypical group and that of POE-CpGs in the typical group as well as the correlation of methylation levels between POE-CpGs, lifestyle and aging were the most associated phenotypic categories for POE-CpGs, in particular for the atypical POE-CpG group (Figure 3b). Smoking status and DNAmTL acceleration were the most associated phenotypes (Figure 3c). After annotating POE-CpGs onto functional regions, a significant enrichment was detected for maternal-smokingexposure-associated atypical POE-CpGs in CpG islands (Figure s2, Table s6). In comparisons between each phenotype’s association with POE-CpGs and its association with the rest of the methylome (non-POE-CpGs), a strong “atypical POE” enrichment was detected for multiple intelligence, aging, parental smoking exposure and lifestyle phenotypes, with verbal intelligence (Mill Hill vocabulary test score) and DNAmTL acceleration displaying the strongest enrichment (Figure 4, Table s7); in contrast, for “typical POE”, only weak enrichments were detected in a few phenotypes (alcohol consumption and maternal smoking exposure)(Figure 4, Table s7).

**Figure 2.**
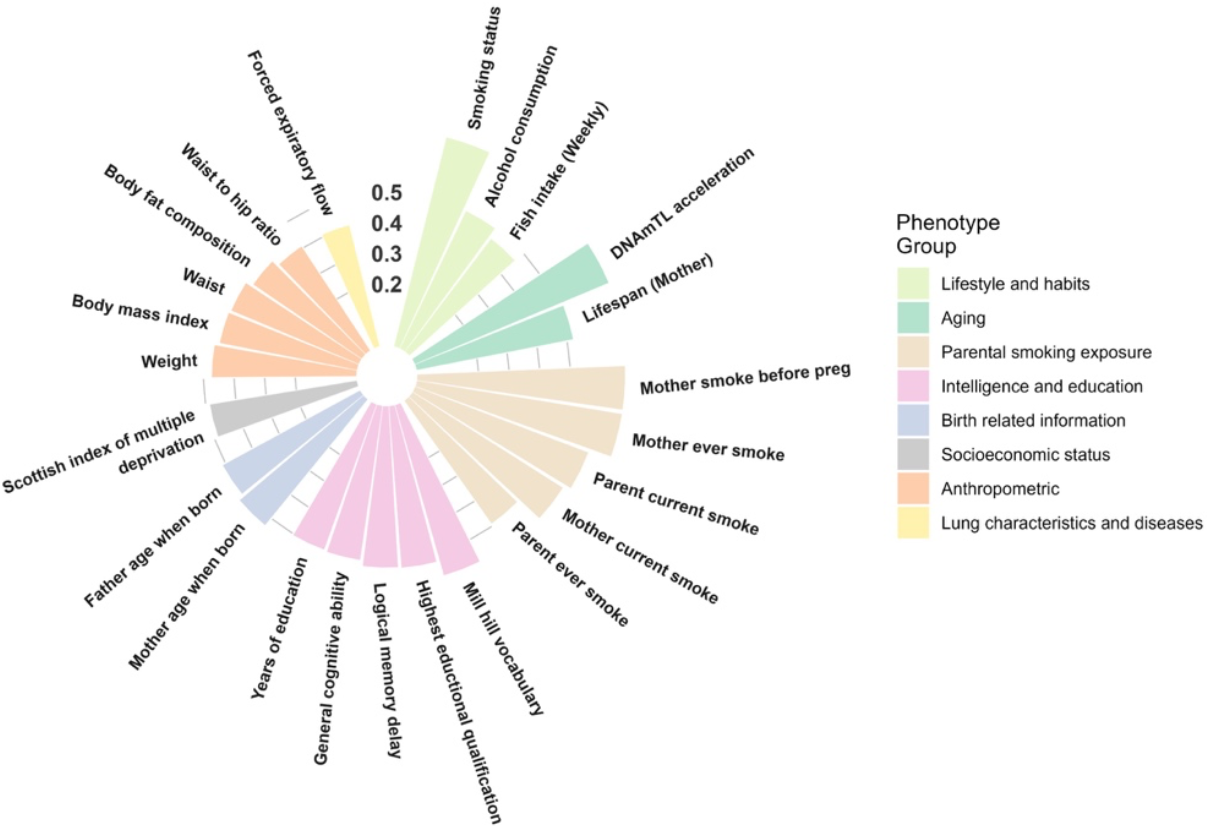
The overall significance of POE-methylation associations for each of the 24 phenotypes associated with at least one POE-CpG. Height of the bar: the average of minus log transformed P values of associations between all POE-CpGs and the given phenotype.

**Figure 3.**
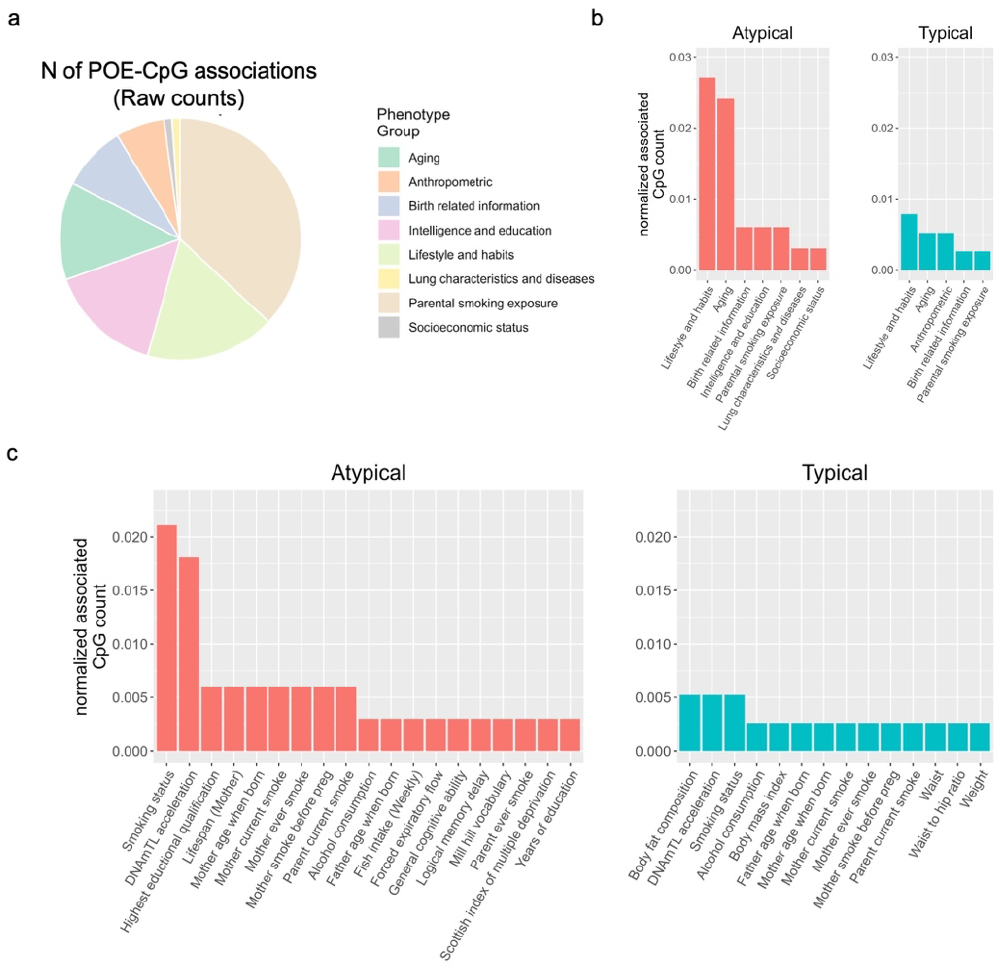
The count of POE-CpGs significantly associated with each phenotype and the category it belonged to. a. Raw counts. b,c: the raw counts were normalized by the total number of POE-CpGs in the atypical or typical group, and the correlation between POE-CpGs(a correlated POE-CpG cluster only counts once). b: normalized counts at categorical level; c: normalized counts at phenotypic level.

**Figure 4.**
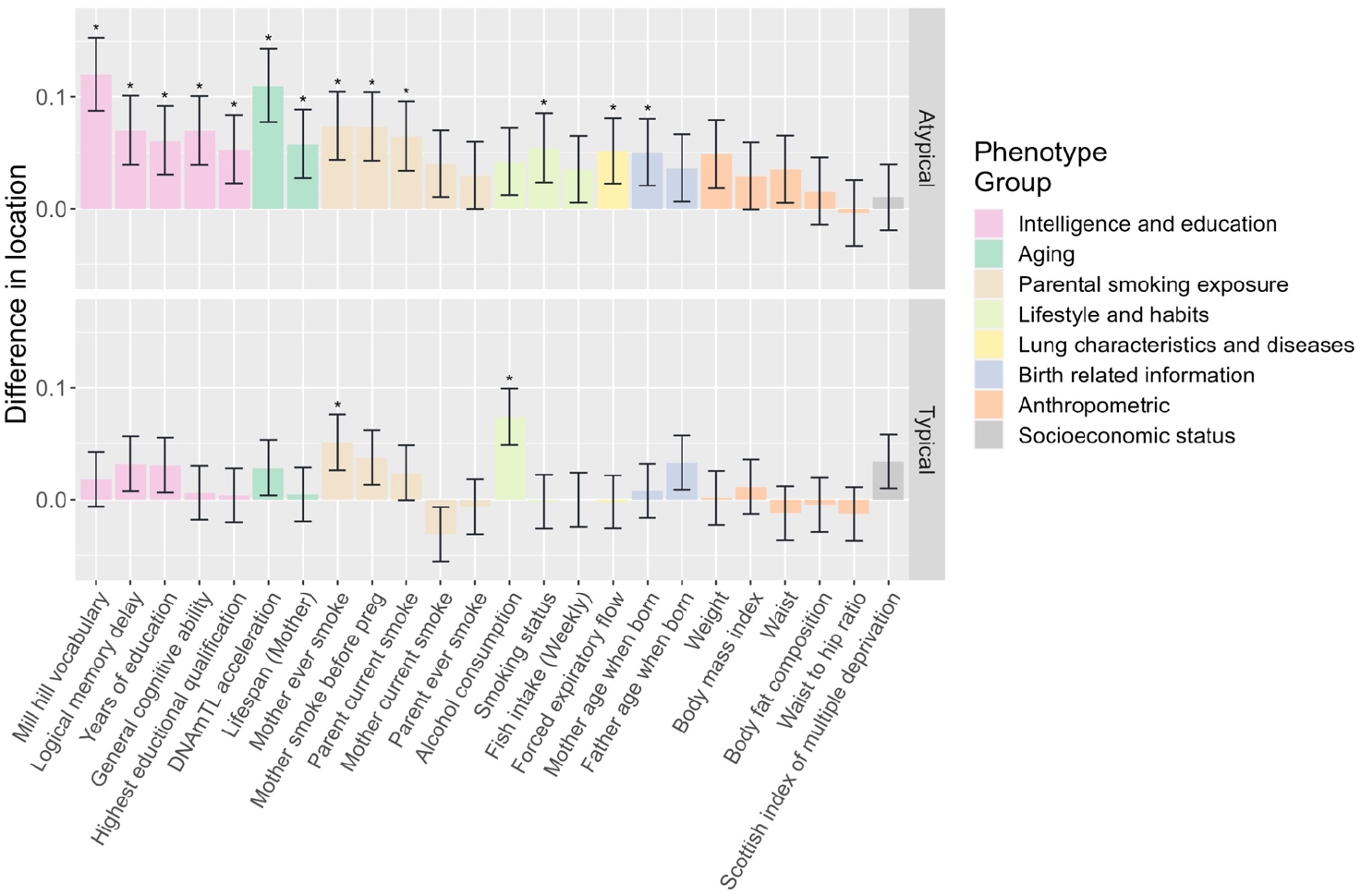
Wilcox test results for comparisons between the methylation association signals from POE regions and the signals from non-POE regions for each of the 24 phenotypes associated with at least one POE-CpG. . *: significant “POE” enrichment was detected on the phenotype (adjusted P < 0.05)

The 92 high confidence associations included cases where a single POE-CpG was associated with phenotypes from multiple trait categories (Figure 5, Table s4). For example, hypermethylation of cg14391737, a POE-CpG located in a CpG shore and an intron of gene *PRSS23(Serine* Protease 23), was simultaneously associated with decreased smoking exposure(self), higher DNAmTL acceleration (longer age-adjusted DNAm telomere length), higher education, higher forced expiratory flow (better lung function) and higher Scottish index of multiple deprivation score (SIMD) (better socioeconomic status). For three POE-CpGs (cg04180046, cg19089201, and cg12803068) in gene *MYO1G* (Myosin IG), hypermethylation was associated with increased maternal smoking exposure, increased smoking exposure(self), and lower intellectual/educational level. There were also cases where a single POE-CpG was associated with multiple phenotypes within a same phenotypic category (Figure 5, Table s4). Multiple POE-CpGs in gene *PRR25* were associated with both maternal and paternal ages when the offspring was born. A POE-CpG in gene *DNTBP1* was associated with anthropometric traits (body fat composition, body mass index, weight and waist). Seven POE-CpGs in gene *CYP1A1/CYP1A2* and one POE-CpG in gene *FRMD4A* were associated with maternal smoking exposures (both current and before pregnancy). Conditional analyses indicated that the multiple associations of these POE-CpGs were not driven by the socioeconomic status (measured as SIMD)(Table s8).

**Figure 5.**
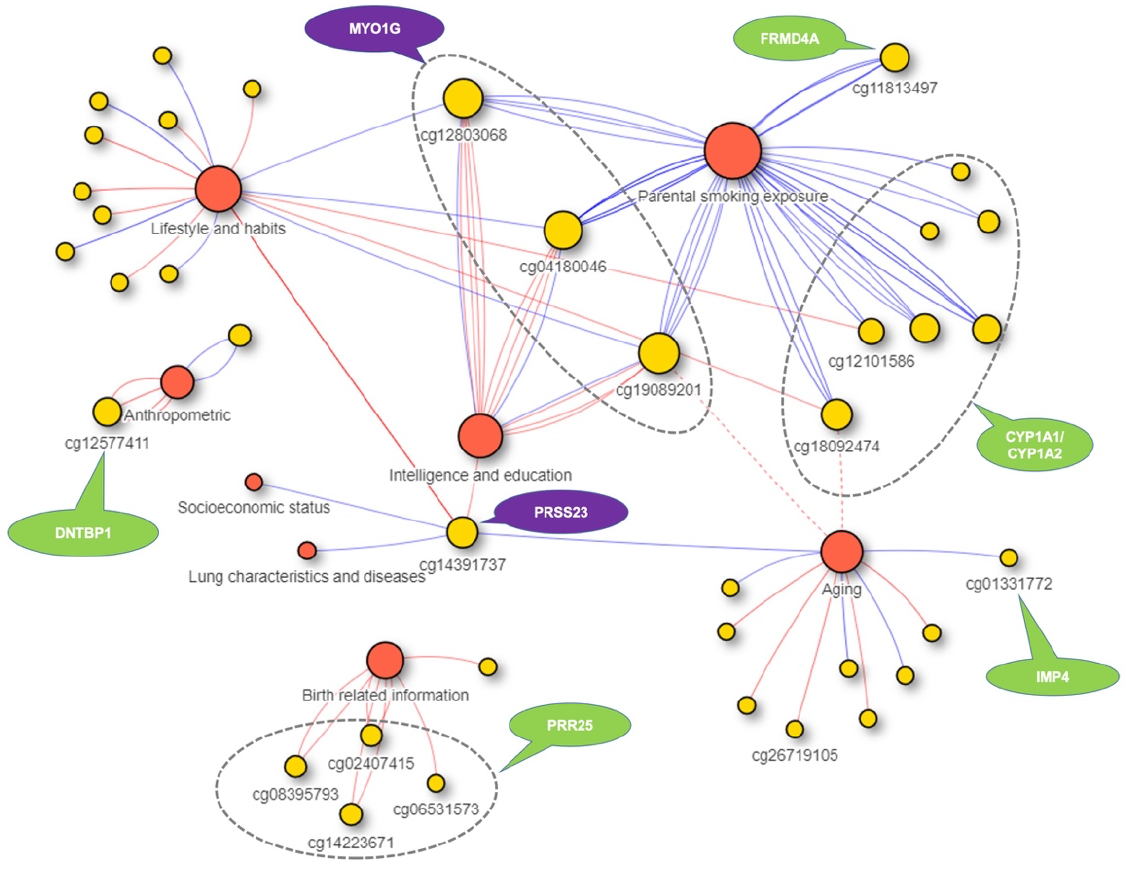
The 92 high confidence associations between POE-CpGs and phenotypes. Red circles: phenotypic categories. Yellow circles: POE-CpG. Each lines represents a significant pair, with red and blue lines representing negative and positive correlation, respectively; the two red dotted lines represent that although statistically significant the associations between cg18092474/cg19089201 with mother’s lifespan were likely to be introduced by association between maternal smoking and mother’s lifespan.

### Atypical POE-CpGs synchronized as co-methylation modules which were associated with aging

We next hypothesized that POE-CpGs could be associated with phenotypes through comethylation networks. To test this, we identified co-methylation modules for atypical and typical POE-CpG groups, respectively. Co-methylation modules were initially constructed in discovery and replication datasets independently, after which “consistent modules” across datasets were identified (methods). The results showed that co-methylation networks were highly reproducible across discovery and replication datasets (Figure s3, Table s9). Six and eight “consistent modules” were identified for atypical and typical POE-CpG groups, respectively (Table s10), and these were used in downstream association tests.

PCs of constituent CpGs’ methylation level were calculated for each “consistent module” for each participant and were used in phenome-wide association tests (methods). Using the discovery dataset, 30 and 2 significant module-PC-phenotype associations were identified (FDR < 0.05) for atypical and typical POE-CpG modules, respectively. Using the replication dataset, 23 (77%) and 2 (100%) of the significant associations were statistically replicated for atypical and typical POE-CpG groups, respectively (Table s11, Figure 6). For the atypical POE group, multiple co-methylation modules were associated with aging phenotypes (DNAmTL acceleration and PhenoAge acceleration) and smoking status; other associations involved intelligence/education traits and maternal smoking exposures (Figure 6). For the typical POE group, weak associations were detected with intelligence/education phenotypes (Figure 6).

**Figure 6.**
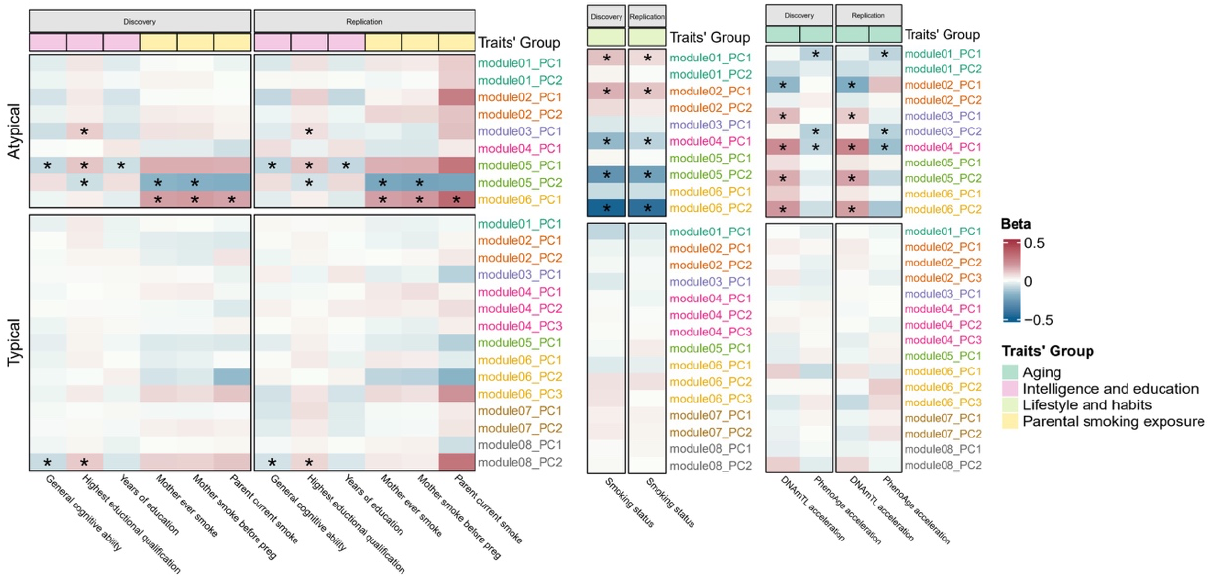
Associations between POE co-methylation module PCs and phenotypes. Only phenotypes with at least one significant result are shown. *: replicated associations.

### An aging-associated atypical POE co-methylation network (module) whose internal methylation connectivity increased with age

POE co-methylation modules’ association with aging implicated that POE-CpGs could associate with aging in an inter-connected and synchronized way. Would the internal methylation connectivity within aging-associated modules alter during aging? To address this, we stratified samples into different age groups (Table s2). For each aging-associated POE co-methylation module, the mean of the methylation connectivity across constituent CpGs was calculated in each age group and compared across groups. The results revealed that the mean of the methylation connectivity of atypical POE module 3 progressively increased with age (Figure 7a, Table s12). The variance of the connectivity of the same module also increased with age, suggesting existence of subgroups (Figure 7a, Table s13).

**Figure 7.**
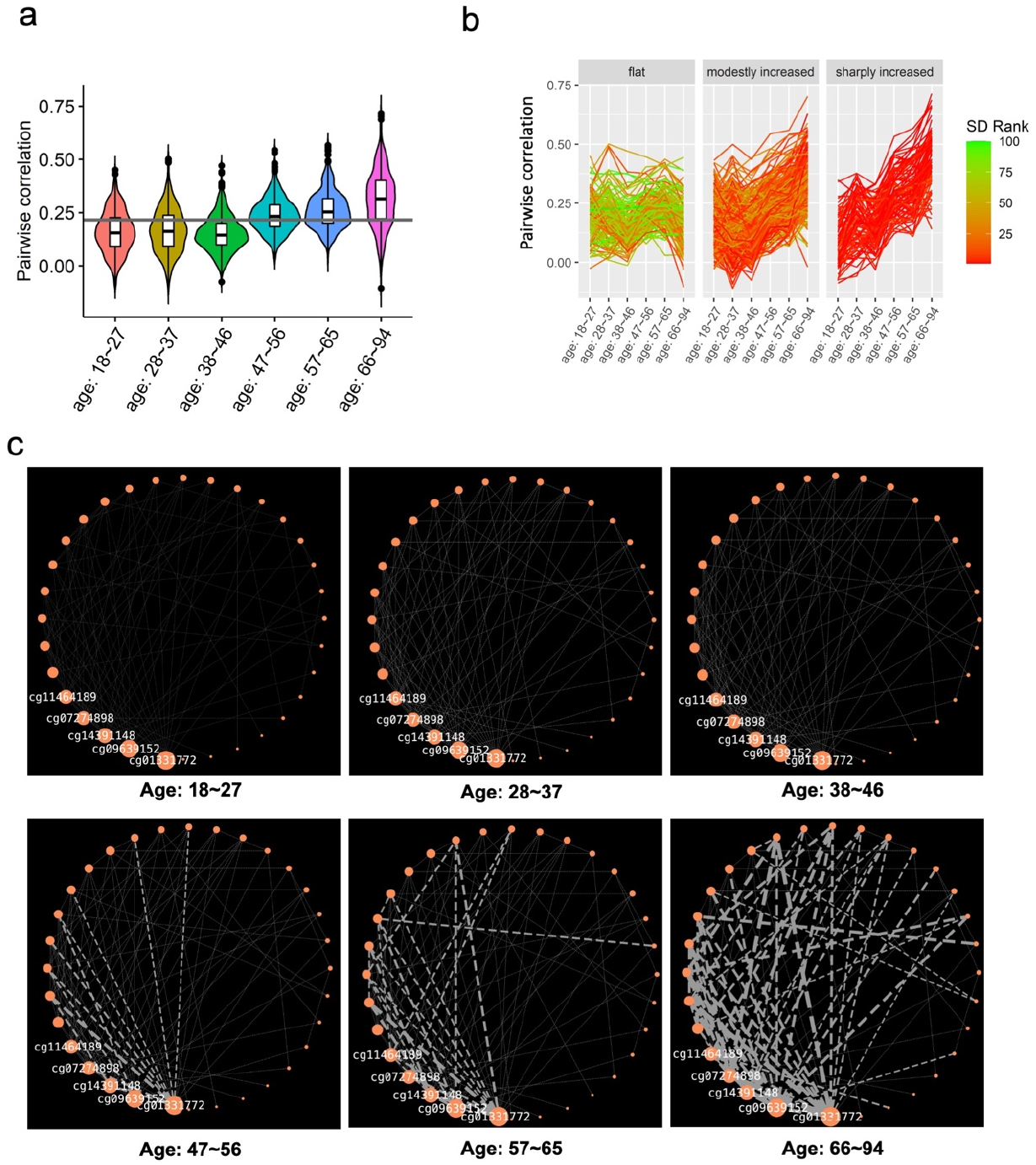
Atypical POE module 3, the module whose internal connectivity increased with age. a. Pairwise between-CpG correlation of constituent CpGs of atypical POE module 3 across different age groups; vertical line: the mean of the methylation correlation across all age groups. b.Three subclusters identified within atypical POE module 3 based on longitudinal trajectory of module connectivity; each line connected the methylation correlation value of a pair of POE-CpGs in different age groups; the color of the line corresponded to the rank of the standard deviation based on the connectivity of POE-CpG pairs across different age groups. c. Methylation connectivity of POE-CpG pairs belonging to the ‘sharp increasing cluster’ in each age group; Orange nodes represented POE-CpGs, the size of orange nodes was scaled by degree centrality (the IDs of the top 5 hub CpGs are shown), the width of edges was scaled by pairwise correlation in samples from each age group, only edges connecting CpGs pairs with an absolute value of correlation larger than 0.4 are shown.

Indeed, based on the longitudinal trajectory of within-module methylation connectivity, our clustering analyses revealed that co-methylated CpG pairs within this module could be further divided into three clusters: a relatively flat cluster(c1), a modestly increasing cluster (c2), and a sharply increasing cluster (c3) (Figure 7b). Within the sharply increasing cluster (c3), five hub CpGs (cg01331772, cg09639152, cg14391148, cg07274898, cg11464189) were identified for displaying the highest centrality, connecting with the largest number of CpGs (Table s14). Also in the sharply increasing cluster (c3), the strength of methylation connectivity, in particular for connections radiating from the five hub CpGs, progressively increased during aging, with the strongest connectivity reached at the oldest (66y-94y) age group (Figure 7c, Table s15). In contrast, none of other constituent CpGs of this module displayed such significant alteration of connectivity strength with age (Table s15). These results revealed the central role of the five hub CpGs in driving the increased methylation connectivity pattern of atypical POE module 3 during aging, suggesting the increased importance of the module and the five hub CpGs at old age groups.

Although the five hub CpGs are in different chromosomes (Figure s4), they are all located within functional regulatory regions as reported by the Roadmap Epigenomics project (58): cg01331772 and cg07274898 are located in promoters (active TSS); cg09639152, cg14391148 and cg11464189 are located in enhancers (mostly bivalent enhancers)(Figure s4, Table s16), implicating the potential of their methylation variation in influencing the expression of nearby genes. Intriguingly, four of the five hub CpGs (Table s16) are located within or nearby a gene encoding a protein that physically interacts with the amyloid beta (A4) precursor protein (APP) in vitro (6.3 fold enrichment, P_fisher_= 5.6×10^-4^)(59).

Among the four hub CpGs located within/near the APP-interactive genes, cg01331772 displayed the highest centrality (Figure 7c, Table s14) and the strongest elevation of methylation connectivity in the sharply increasing cluster (c3) in the comparison between youngest and oldest age groups (Table s15). This CpG is located in a promoter, and is 987 bp downstream of gene *CCDC115* (Coiled-Coil Domain Containing 115) and 4791 bp upstream of gene *IMP4* (IMP U3 Small Nucleolar Ribonucleoprotein 4) (Figure s4). In blood, the methylation level of this CpG was positively associated with the mRNA expression of *IMP4* (P_eQTM_=9.7*10^-7^), as reported by a recent eQTM (expression quantitative trait methylation) study(60). In brain, the methylation levels of cg01331772 and the mRNA expression of *IMP4* were genetically positively correlated in our OMIC-based SMR analysis (Beta_SMR_=0.35, P_SMR_adjusted_=8.4×10^-4^, P_HEIDI_unadjusted_=0.1). For this CpG, the brain-blood methylation correlation was relatively high (Rho=0.54, P=0.01, Figure s5)(61), suggesting that methylation of cg01331772 in blood could be indicative for expression of *IMP4* in brain tissues. Notably, *IMP4’s* mRNA expression decreased significantly in Alzheimer’s disease (AD) patients as compared to controls in both temporal cortex (P=0.003) and prefrontal cortex (P=2.6×10^-6^), the two most relevant brain regions for AD pathogenesis (Figure s6)(62–64). Putting these observations together, increased methylation of the hub CpG cg01331772 in blood may imply higher expression of *IMP4* in AD-susceptible brain tissues, which can be protective for AD.

Interestingly, the associations between the hub CpG cg01331772 and aging dramatically changed cross different life stages. The PC1 of the atypical POE module 3, explaining 28.7% of the methylation variance within that module and having a positive loading from cg01331772 (Table s17), displayed a similar association pattern with aging. In brief, using samples from the full age spectrum in GS:SFHS (18-94y), at single-CpG level, the hypermethylation of cg01331772 was associated with older chronological age (Table s18) and longer age-adjusted DNAmTL (higher DNAmTL acceleration. Table s4); at modular level, the PC1 of atypical POE module 3 displayed similar association patterns (Table s11). Why would the methylation of cg01331772 and the PC1 of atypical POE module 3 increase with chronological age while displaying a positive association with DNAmTL acceleration at the same time? The seemingly contradictory observations were disentangled by our age-stratified analyses. We found that starting with the youngest adult years (18-27y), the methylation of cg01331772 significantly increased with age, but the slope decreased to an insignificant level after the middle age was reached (Figure 8). In contrast, no association between cg01331772 and DNAm-predicted telomere length was observed until middle age, after which a positive association started to arise and became much stronger in older age groups (Figure 8). As a consequence, a significant interactive effect between chronological age and cg01331772’s methylation effect on DNAmTL acceleration was detected (P_interaction_=2.2 x 10^-8^), whereby the methylation of this CpG only manifesting significant positive association with DNAmTL acceleration in old age groups (Figure 8). Similar association patterns were observed at the PC1 of atypical POE module 3 (Figure 8). These combined results revealed the importance and the complexity of the role of POE comethylation networks and their hub POE-CpGs in human aging.

**Figure 8.**
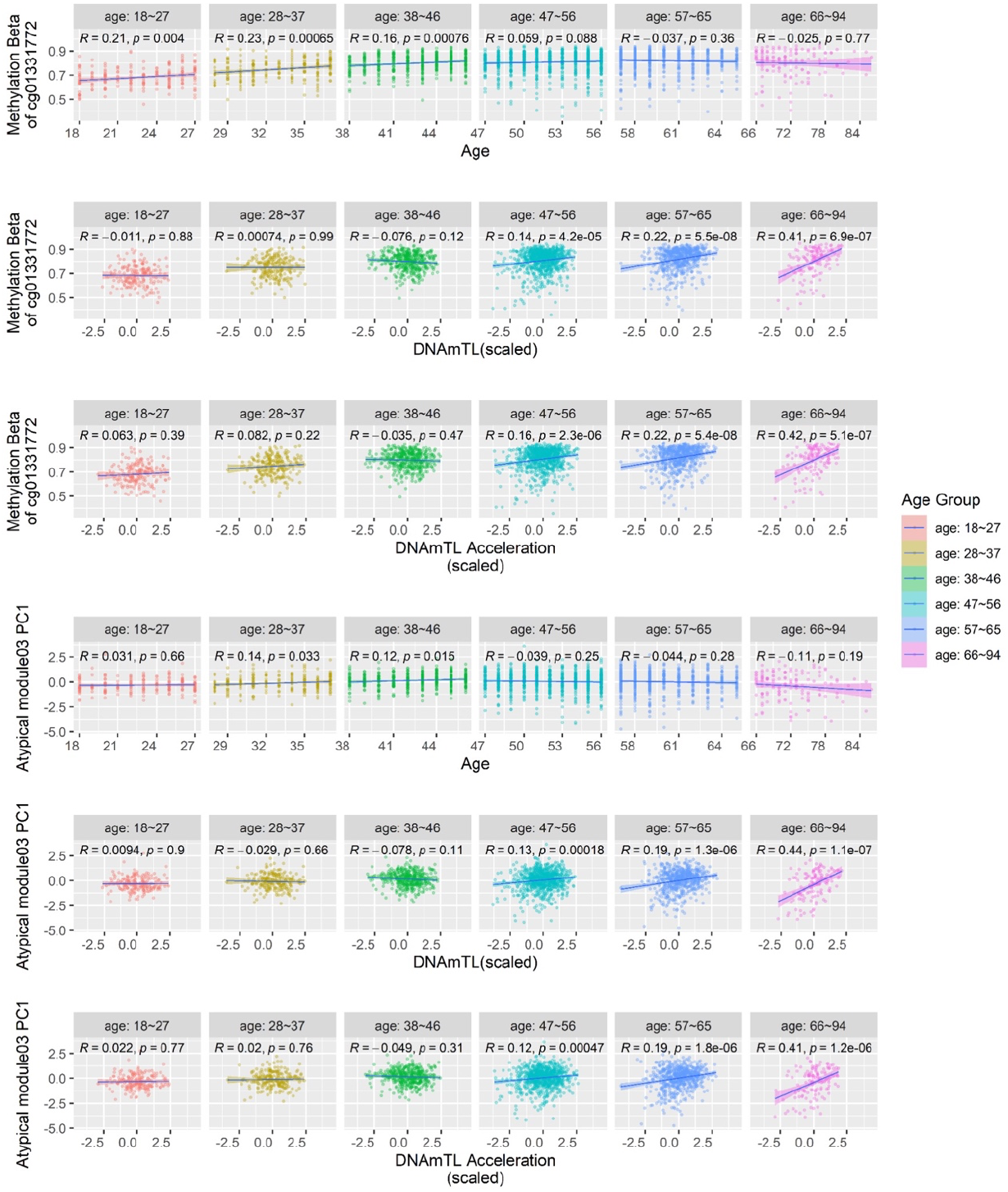
The atypical POE module 3 and its hub POE-CpG cg01331772’s association with age, DNAmTL and DNAmTL acceleration in different age groups.

### High levels of methylation heterogeneity and increased epigenetic drift (information loss with age) of atypical POE-CpGs

As mentioned above, both single-CpG- and network-based analyses supported the special link between POE-CpGs (atypical group in particular) and aging. We next examined whether those CpGs manifested additional aging-related features. In the DNA methylation context, Shannon entropy measures the level of methylation heterogeneity: the higher the Shannon entropy is, the higher the heterogeneity is, the less predictable the methylation condition in a cell population is (52, 65, 66). Shannon entropy is maximized at intermediate methylation levels (Beta=0.5) and minimized in extreme methylation levels (Beta=0 or 1). It has been known that aging was accompanied with an increased epigenetic drift (the loss of information stored in the epigenome), reflected as the age-related increment of average methylation Shannon entropy in the epigenome as a whole, or in a few aging-related functional CpG sets with a faster drift rate(52, 65, 66). Here, we compared the Shannon entropy for POE-CpGs, in particular for those belonging to the atypical group, to epigenetic clock CpGs and the rest of epigenome.

The results showed that taking POE-CpGs as a whole, their Shannon entropy was significantly higher than the global level of the methylome, higher than Horvath clock CpGs and Hannum clock CpGs and slightly lower than DNAmTL clock CpGs (Figure 9a, Table s19). After we stratified POE-CpGs into subgroups, the atypical POE group’s Shannon entropy was significantly higher than that of the typical group; the aging-associated POE-CpGs displayed higher Shannon entropy than the POE-CpGs without an association with aging (Figure 9a, Table s19). In terms of epigenetic drift (information loss) with age, Shannon entropy of all CpGs groups significantly increased with age, with the atypical POE-CpG group displaying faster information loss with age as compared to the typical POE-CpG group and the global methylome (Figure 9b, Table s20).

**Figure 9.**
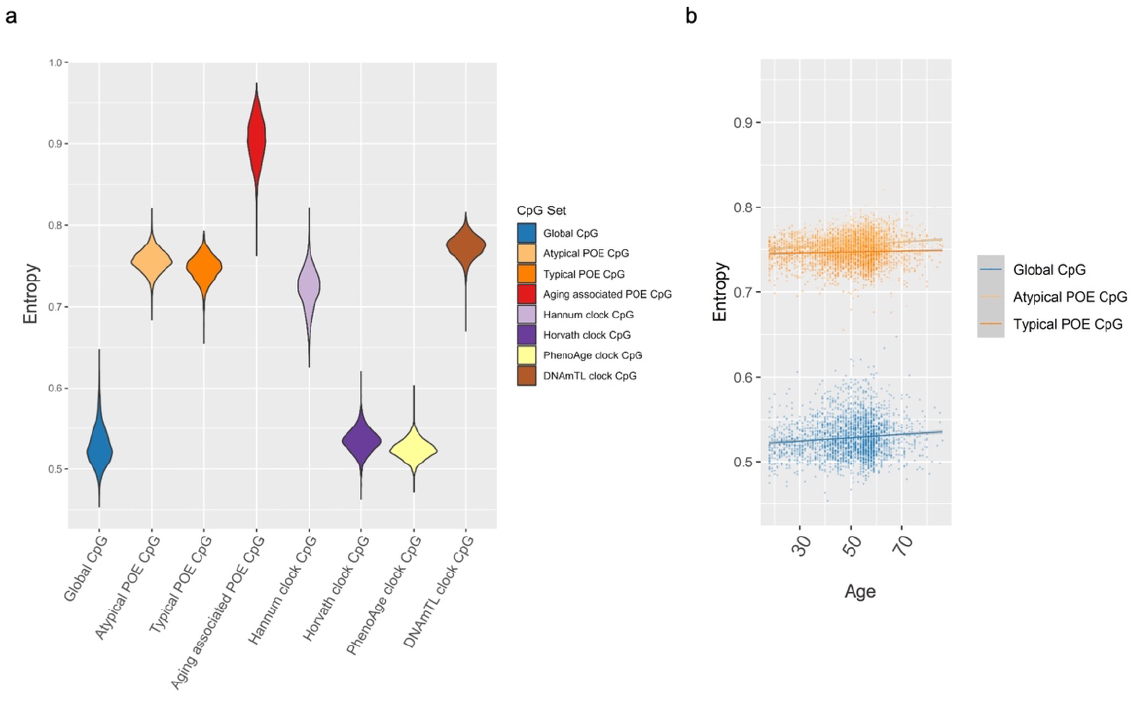
Shannon entropy of POE-CpGs and CpGs from other categories.

### Intrinsic connection between clock CpGs and atypical POE-CpGs

Given the shared high Shannon entropy feature between POE-CpGs and clock CpGs, we wondered whether the POE-CpGs and clock CpGs are intrinsically connected. To address this, circular permutation was applied to test whether atypical/typical POE-CpGs were more correlated with clock CpGs compared against randomly selected CpG sets of the same size from the methylome. The results revealed a significantly higher correlation between atypical POE-CpGs and constituent CpGs of all of the four popular clocks when compared to the random CpG sets, whereas this was not observed in the typical POE-CpG group (Figure 10).

**Figure 10.**
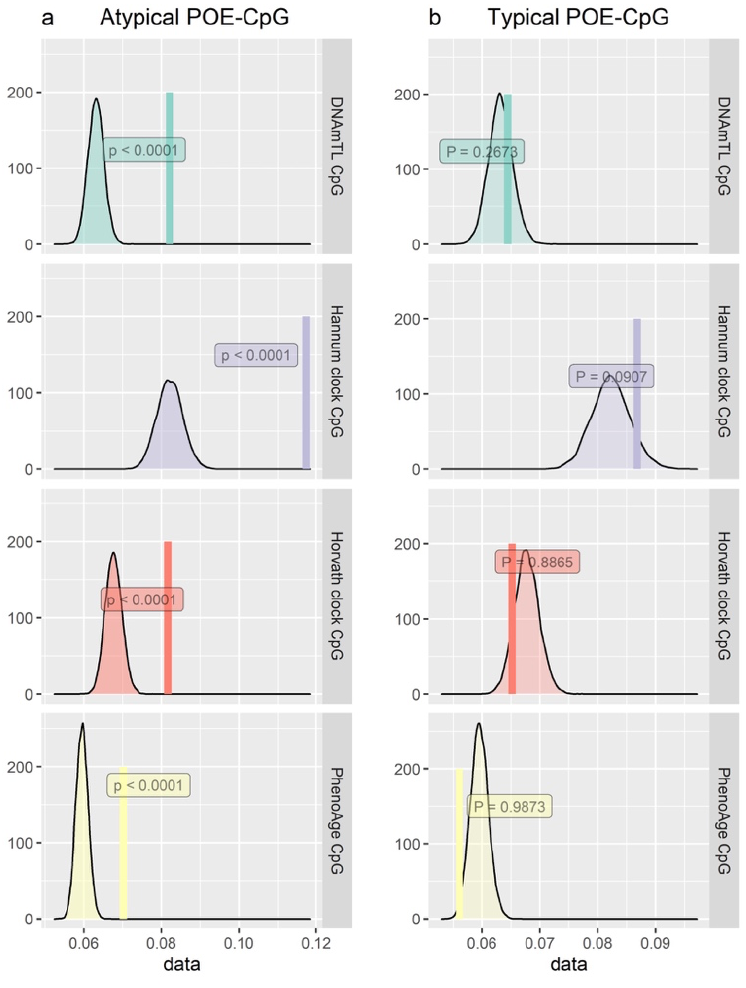
Permuted (the distribution) and observed correlation (the vertical line) between POE-CpGs and constituent CpGs of epigenetic clocks.

## Discussion

In this study, we systematically examined associations of POE-influenced methylome (POE-CpGs) with adult aging, early/late environmental exposures, and health-related phenotypes. Single-CpG-based analyses revealed replicated and enriched methylation associations with lifestyle (smoking status), aging (DNAmTL acceleration), parental (maternal) smoking exposure, and intelligence phenotypes in atypical POE-influenced regions. Co-methylation analyses indicated that at least a proportion of atypical POE-CpGs were associated with these phenotypes in a modularized way. We additionally reported the age-related increment of internal connectivity in an aging-associated atypical POE co-methylation module. For that module, we identified the hub POE-CpGs that likely drive the increment of connectivity, and uncovered dynamic aging-association patterns of the module and its top hub CpG across different life stages. Finally, compared to the rest of methylome, atypical POE-CpGs displayed high levels of methylation heterogeneity, fast information loss with age, and high methylation correlation with clock CpGs, which further provided evidence for the special link between atypical-POE-influenced methylome and human aging.

At single-CpG level, we found that atypical POE-influenced methylome was sensitive to both early life factors such as maternal smoking exposure and parental age when the offspring was born, and later life exposures such as smoking and alcohol consumption. Meanwhile, atypical POE-CpGs were also strongly associated with aging and health-related phenotypes such as intelligence in adulthood (Figure 3, Figure 4). Importantly, we detected cases where the same single POE-CpG was simultaneously associated with both environmental exposure (such as maternal smoking exposure or lifestyle), adult aging and/or with health-related phenotypes (such as intelligence). Our observation of the associations between cg14391737, an intronic POE-CpG located of gene *PRSS23*, with smoking status and forced expiratory flow (Figure 5), was in line with previous MWAS papers that identified cg14391737 as a smoking- and lung cancer-associated CpG (67, 68). Here, we uncovered its additional associations with education, socioeconomic status and DNAmTL acceleration. Our observation that multiple CpGs within the gene *MYO1G* were associated with maternal smoking exposure, smoking status and the highest educational qualification, was consistent with previous studies (69–71). Importantly, we uncovered the new associations of those early and late-environmental-sensitive CpGs in *MYO1G* with multiple intelligence measurements in adults (Table s4). These results supported well our hypothesis that POE-influenced epigenome could act as a hub position in the interplay of early/late life exposures, adult health and adult aging.

At network level, we found that the methylation levels of a proportion of POE-CpGs fluctuated jointly as co-methylation modules (both in *cis* and *trans*). Consistent with results from single-CpG-based analyses, module-level methylation-phenotype associations revealed the association of the shared methylation variation of multiple atypical POE-CpGs with aging, smoking, maternal smoking exposure, and intelligence. These results suggested that early and late environment may influence atypical POE-CpGs in groups rather than individually, and that aging-associated atypical POE-CpGs can function a modularized way. The aging-associated POE co-methylation networks were not stable throughout the life. We found that atypical POE module 3, one of the aging-associated POE-co-methylation modules, displayed increased connectivity when humans get older (Figure 7). Five hub POE-CpGs were identified for their central role in driving this change, and intriguingly, the majority of the hub POE-CpGs appeared to link to APP-interacting proteins. In particular, the module centrality and aging-associated connectivity change were most prominent in cg01331772, a promoter CpG that was likely capable of regulating expression of *IMP4*, the gene both interacting with APP and displaying significant downregulation in AD patients in two AD-relevant brain regions (Figure s6). These findings coincided with a previous finding suggesting that at methylome-wide level, the aging-associated co-methylation module was enriched for promoter CpGs located nearby genes downregulated in early disease stage of AD(72). Our results suggested the central role of *IMP4’s* regulatory CpG cg01331772 in POE-related modularized methylation alteration during the aging process.

The complexity of the role of atypical POE module 3 and its hub CpG cg01331772 in human aging can be further revealed by integrating existing evidence with our single-CpG- and network-based results. Previous studies have recognized that methylation of cg01331772 persistently increased at early life stage (age ≤10y)(73–75). Our stratification analyses covered a wide age spectrum of human adults (18-94y) and showed that the age-associated elevation of methylation in this CpG continued until one’s middle age. At older age groups, this CpG was no longer associated with chronological age, but surprisingly, shifted to be associated with increased DNAm-predicted telomere length (DNAmTL) and age-adjusted DNAmTL (DNAmTL acceleration), with the strongest association appearing in the oldest age group (66-94y) (Figure 8). The PC1 of atypical POE module 3 where cg01331772 has a positive loading also followed this pattern (Figure 8). Previously, IMP4 has been reported as a component of telomerase whose function was to maintain/elongate telomere(76, 77). Here, we found that hypermethylation of cg01331772, a likely-regulatory POE-CpG for *IMP4*, was associated with longer predicted-telomere-length at old age groups. Importantly, since cg01331772 acted as a hub CpG for an aging-associated co-methylation module that become highly self-connected at old age, this effect has the potential to propagate through the comethylation network. These observations not only unveiled new targets (cg01331772 and other constituent CpGs of atypical POE module 3) for future biomarker and intervention studies of aging, but also highlighted that, in order to comprehensively evaluate the multiplex role of functional CpGs such as POE-CpGs in human aging, it is necessary to consider the effects both when they act as individual sites and act as constituent members in a network, consider the dynamic association patterns in stratified age groups, and consider both the mean difference and the connectivity difference in the methylation network.

POE-CpGs also manifested other aging-related features such as a high degree of methylation heterogeneity, a fast epigenetic drift with age and a strong methylation correlation (the atypical POE group) with constituent CpGs of four epigenetic clocks. As the clock CpGs were well known for the associations with aging, here, they were used to compare with POE-CpGs to benchmark the aging-related features of those CpGs. Previous studies have reported the general genome-wide trend of loss of methylation information content (manifested as increased entropy) with age (5, 52), the high entropy in epigenetic clock CpGs compared to the rest of genome(51), and the positive association between methylation entropy and age acceleration(5). Here, our study showed that as a specialized group of CpGs, POE-CpGs (atypical group) not only lost methylation information with age in a rate that was faster than the rest of global methylome, but also displayed unusually high methylation heterogeneity (entropy) that was even higher than the constituent CpGs of three popular epigenetic clocks (Horvath, Hannum, PhenoAge). POE-CpGs’ entropy was only slightly lower than that of DNAmTL CpGs, but was higher when only considering aging-associated POE-CpGs. The high entropy feature shared between POE-CpGs and clock CpGs inspired us to hypothesize that POE-CpGs and clock CpGs were intrinsically interconnected, given their shared association with aging. Indeed, although there were only 10 CpGs labeled as both POE-CpGs and clock CpGs, we found a much higher correlation between atypical POE-CpGs and clock CpGs for all of the four clocks tested compared to the correlation with the rest of the methylome (Figure 10). This was not observed in typical POE-CpG group, consistent with our observation that the atypical POE-CpG group displayed much stronger and enriched associations with aging phenotypes compared to typical POE-CpGs (Figure 3, Figure 4). It is noteworthy that clock CpGs have been known for their ability to predict aging, whereas POE-CpGs were identified for the special heritable pattern introduced by early life events (imprinting or early environmental effect); the shared features between the two classes of CpGs further supported the association between atypical POE-CpGs and aging.

There are limitations in this study. First, the associations we reported were discovered and replicated in a Scottish population, future studies are needed to replicate our findings in other populations. Second, survival bias could influence the estimates of the methylation connectivity for the atypical POE module 3 in old age groups, given the cross-sectional feature of our samples. Although the methylation connectivity of that module has already started to increase at young age (Figure 7), which suggests that the overall increment trend is less likely to suffer from the survival bias issue, future longitudinal data will help to validate our findings in old age groups. Third, aging-associated methylation dynamics can be confounded by varied cell proportions of rare cell types. Although we accounted for cell count effects by pre-adjusting or jointly fitting estimated proportion of major blood cell types as covariates, the proportions of rare cell types can vary substantially across age groups but are difficult to estimate, this could confound methylation analyses using data generated from bulk tissues like ours. Fourth, our analyses on effects from early environmental exposures on POE-CpGs were largely limited to parental smoking. Future analyses using samples with richer and higher resolution records of early environmental exposures would allow a more comprehensive evaluation of effects from early environmental exposures and lifetime consequences. Finally, although longer telomere (and higher telomere length acceleration) in non-tumor tissues is usually considered protective, there is also evidence suggesting that longer telomeres can be associated with higher risk of cancer (78). We suggest that conclusions regarding longevity from our DNAmTL acceleration analyses should be made with caution, future studies that investigating the association between POE-CpGs and longevity directly are warranted.

In conclusion, our phenome-wide human methylation analyses identified strong and enriched association between the atypical-POE-influenced methylome and adult aging, and between atypical-POE-influenced methylome and early/late exposures at both single-CpG and network levels. The shared high epigenetic drift features and the intrinsic connections between atypical POE-CpGs and clock CpGs were also revealed. The identified single POE-CpGs and POE co-methylation modules provided new targets for future biomarker and intervention studies and added novel supporting evidence for the “early development of origin” hypothesis for adult aging.

## Acknowledgements

YZ is supported by National Key Research & Development Program of China (No. 2021ZD0202000) and the General Program of National Natural Science Foundation of China (81971270). AMM is supported by NIH award R01MH124873 and by UKRI award MR/W014386/1. CA, CH, CSH and PN want to acknowledge support from the MRC Human Genetics Unit programme grant, ‘Quantitative traits in health and disease’ (U.MC_UU_00007/10), and grant MC_PC_U127592696. Generation Scotland has received core funding from the Chief Scientist Office of the Scottish Government Health Directorates CZD/16/6 and the Scottish Funding Council HR03006. Genotyping of the GS samples was carried out by the Genetics Core Laboratory at the Wellcome Trust Clinical Research Facility, Edinburgh, Scotland and was funded by the UK MRC and the Wellcome Trust (Wellcome Trust Strategic Award ‘Stratifying Resilience and Depression Longitudinally’ (STRADL) Reference 104036/Z/14/Z). The DNA methylation (DNAm) profiling and analysis was supported by Wellcome Investigator Award 220857/Z/20/Z and Grant 104036/Z/14/Z (PI: AM McIntosh) and through funding from NARSAD (Ref: 27404; awardee: Dr DM Howard) and the Royal College of Physicians of Edinburgh (Sim Fellowship; Awardee: Dr HC Whalley)”. GS:SFHS: we want to acknowledge support from Genetics Core Laboratory at the Wellcome Trust Clinical Research Facility (Edinburgh, Scotland) for genotyping of the GS samples. We are grateful to all the families who took part, the general practitioners, and the Scottish School of Primary Care for their help in recruiting them, and the whole Generation Scotland team, which includes interviewers, computer and laboratory technicians, clerical workers, research scientists, volunteers, managers, receptionists, healthcare assistants and nurses.

## Data sharing statements

Summary statistics supporting the conclusions of this article are included within the article and its additional files. GS:SFHS: Generation Scotland data are available from the MRC IGC Institutional Data Access / Ethics Committee for researchers who meet the criteria for access to confidential data. Generation Scotland data are available to researchers on application to the Generation Scotland Access Committee. The managed access process ensures that approval is granted only to research which comes under the terms of participant consent which does not allow making participant information publicly available.

