## Supplementary materials for "Phenome-wide analysis identifies parent-of-origin effects on the human methylome associated with changes in the rate of aging"

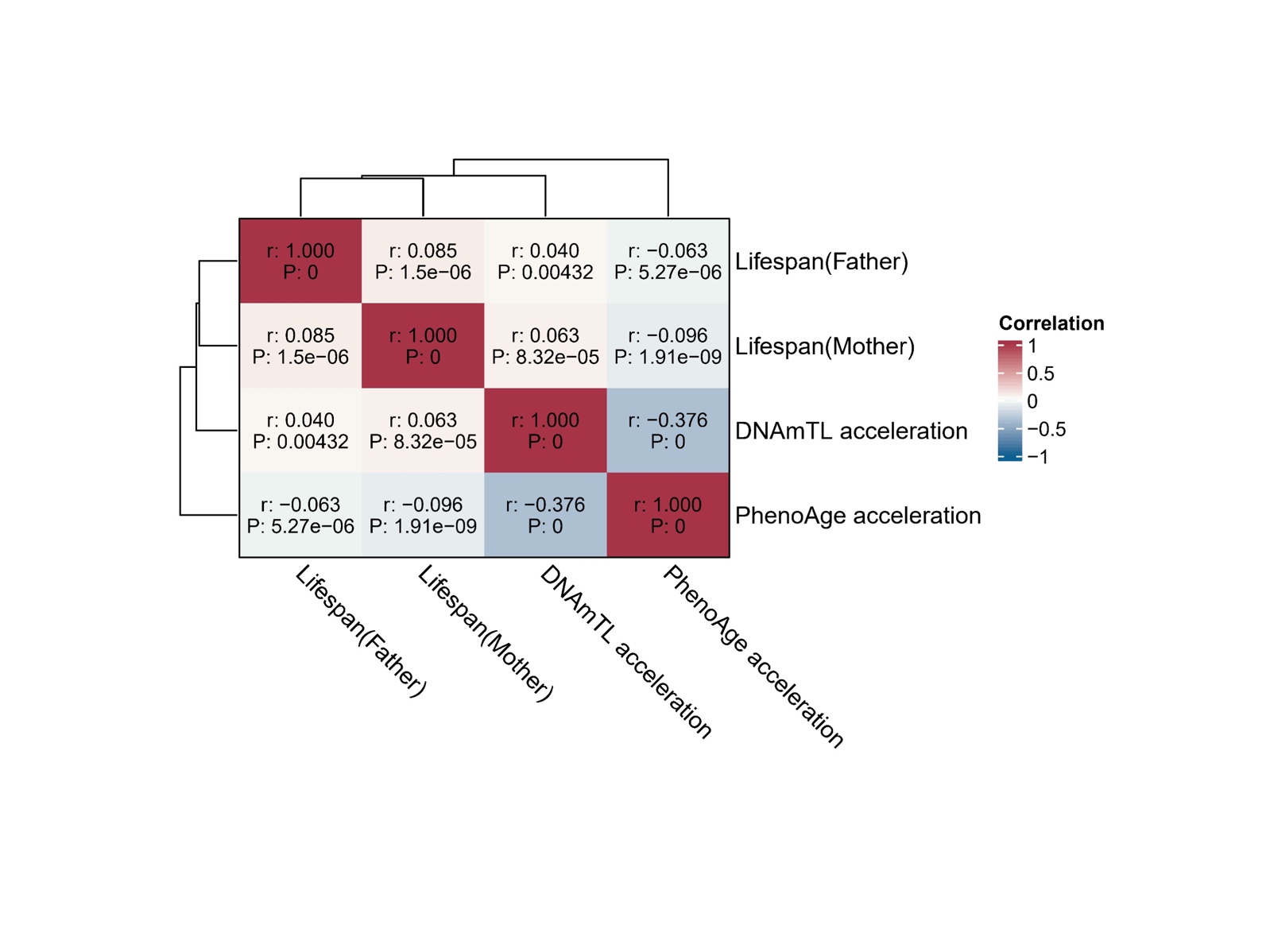

**Figure s1. Phenotypic correlations between the four aging phenotypes.**

\
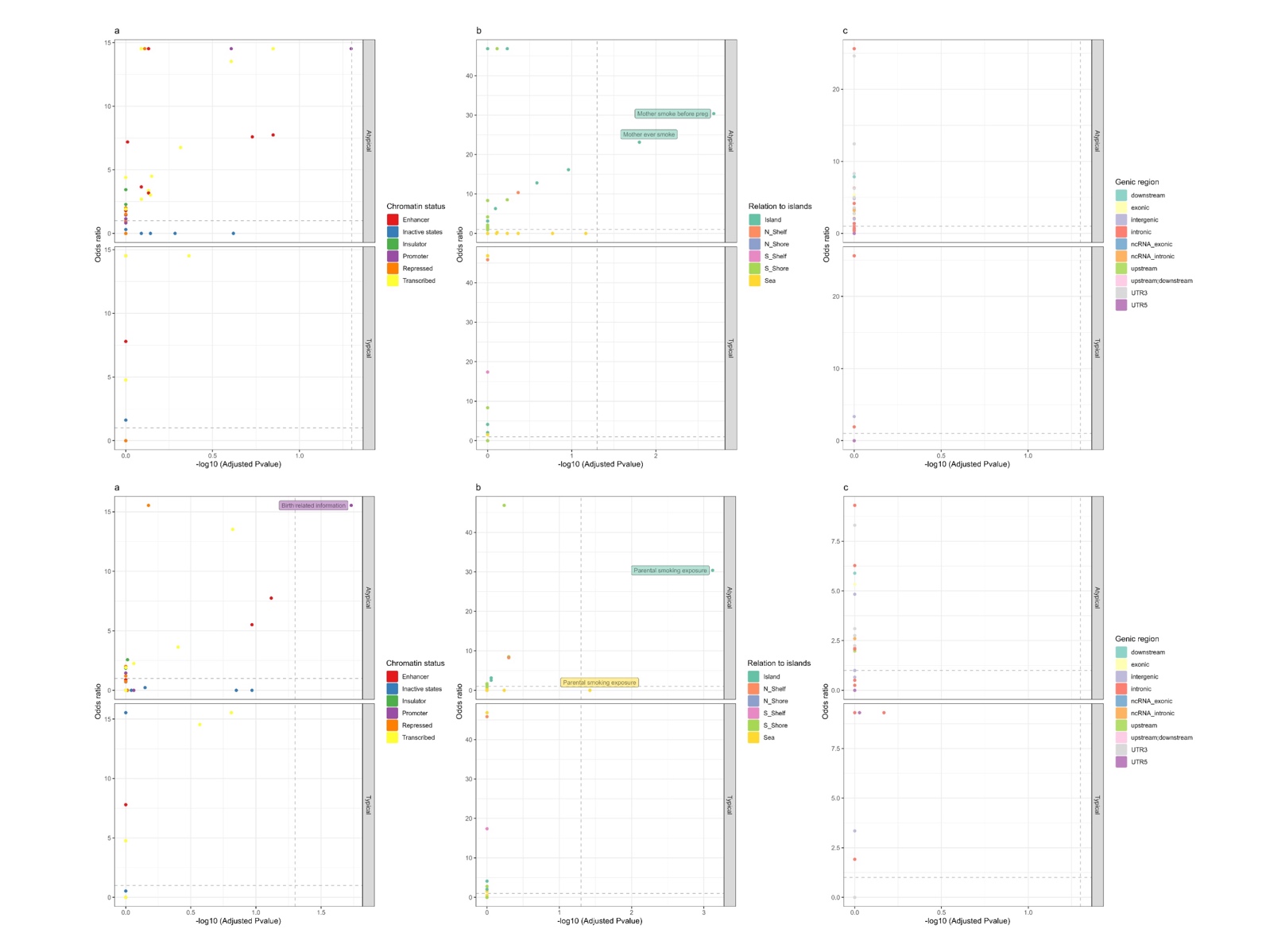

**Figure s2. Functional enrichment of associated POE-CpGs for each phenotype and each phenotypic category.** Upper figures: phenotypic level. Bottom figures: categorical level.

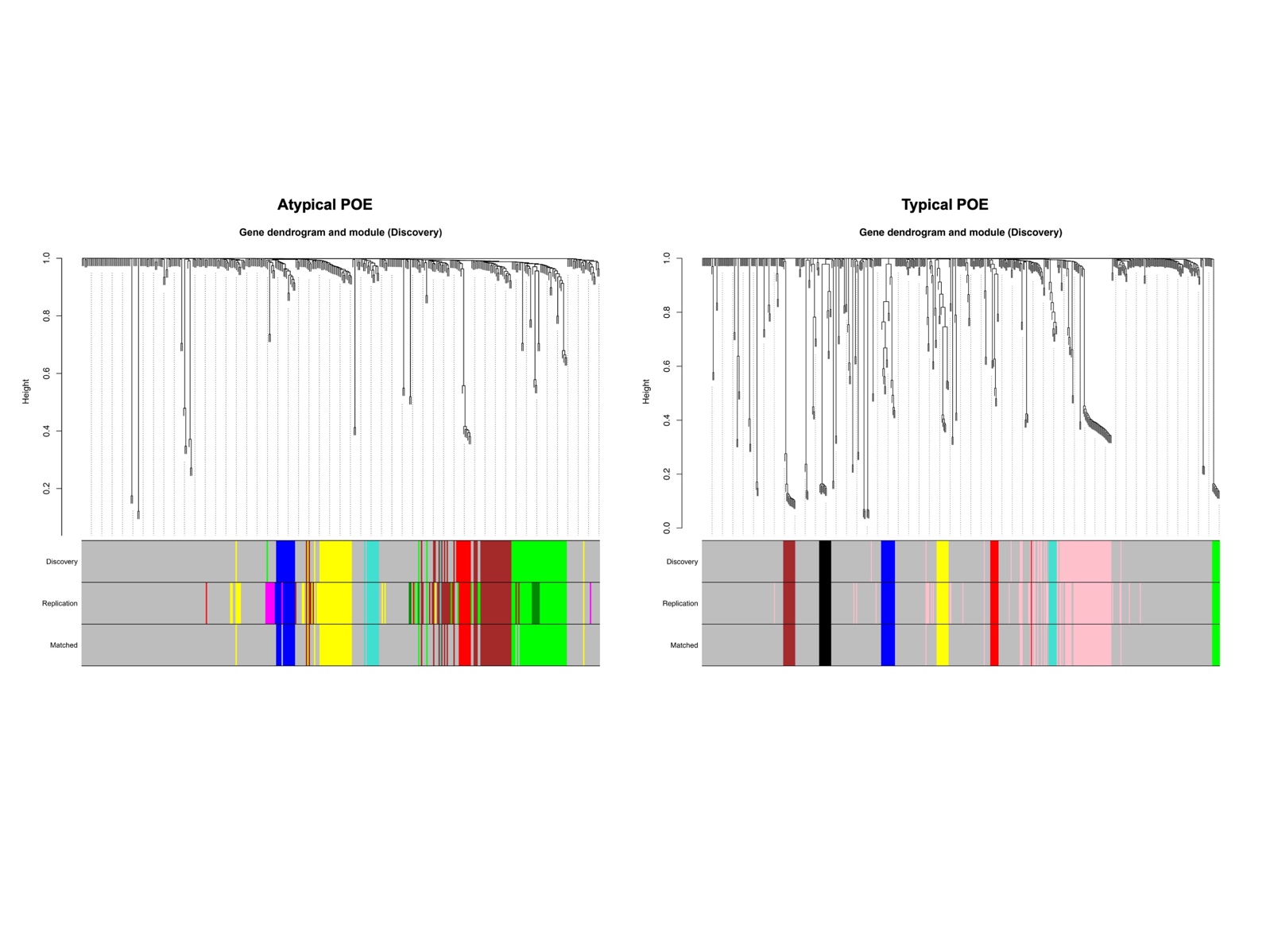

**Figure s3. Identified WGCNA POE co-methylation modules in discovery and replication datasets.**

a.

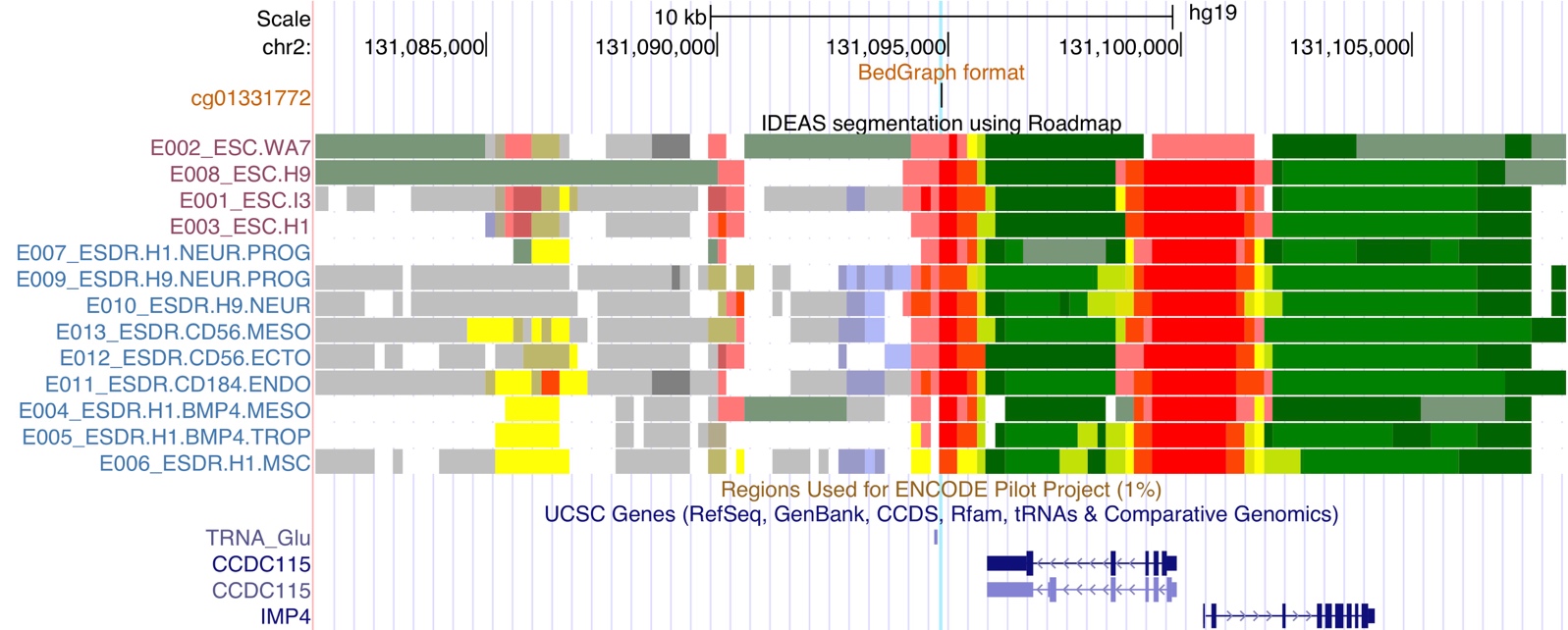

b.

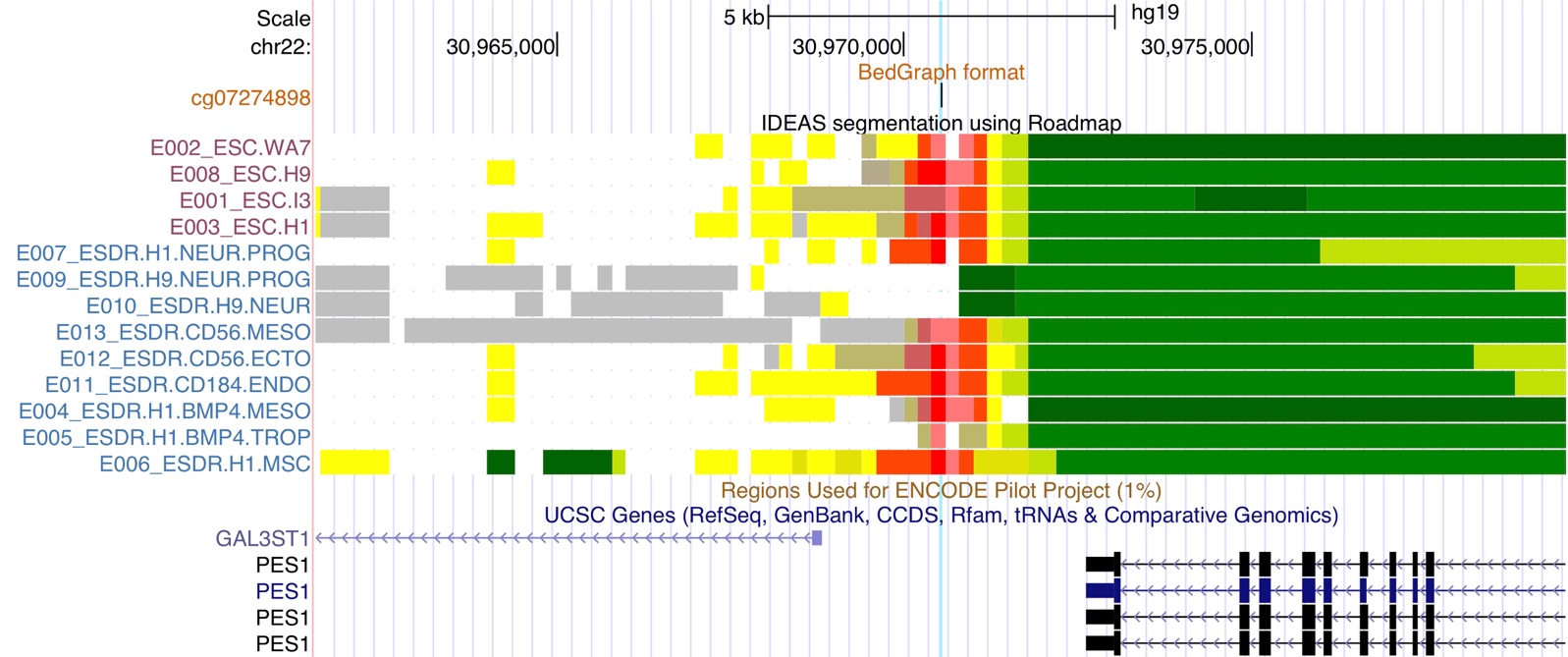

c.

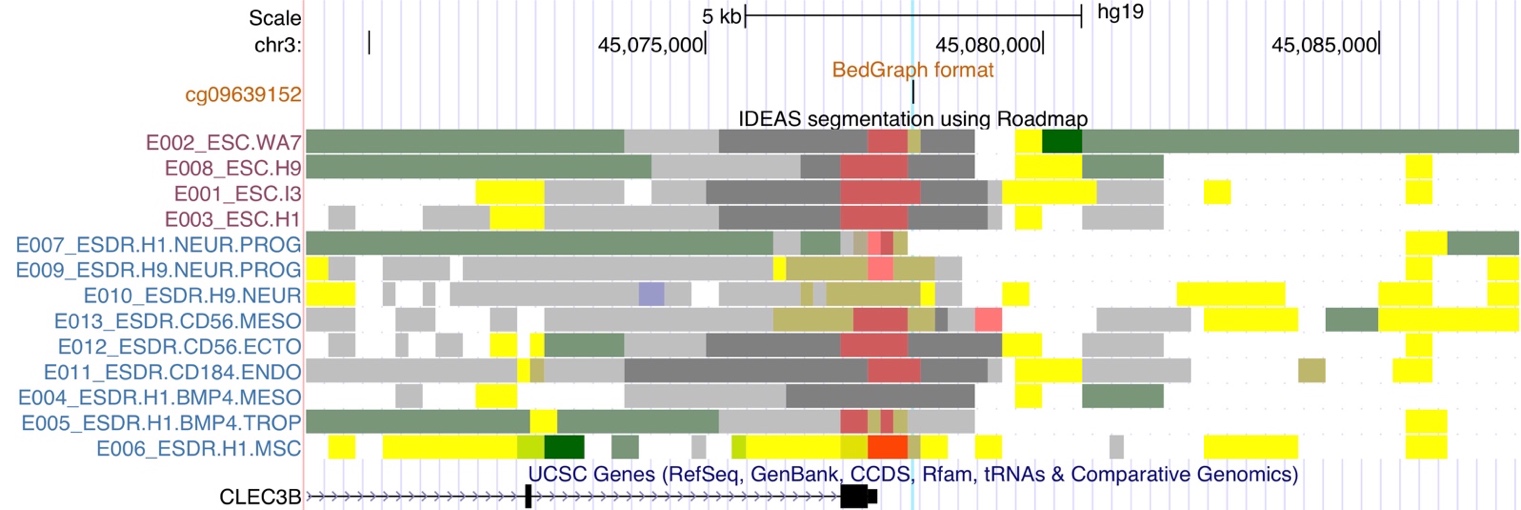

d.

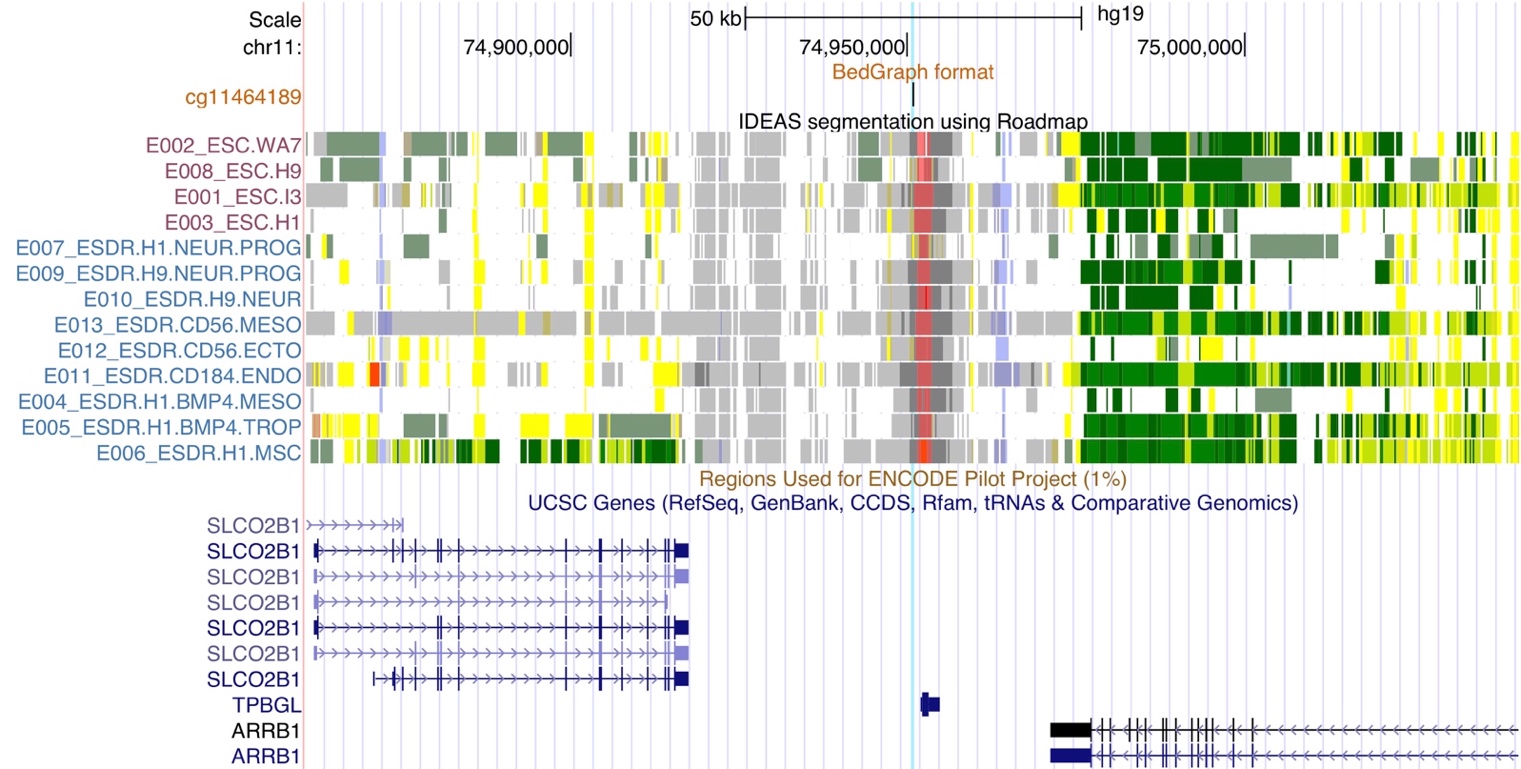

e.

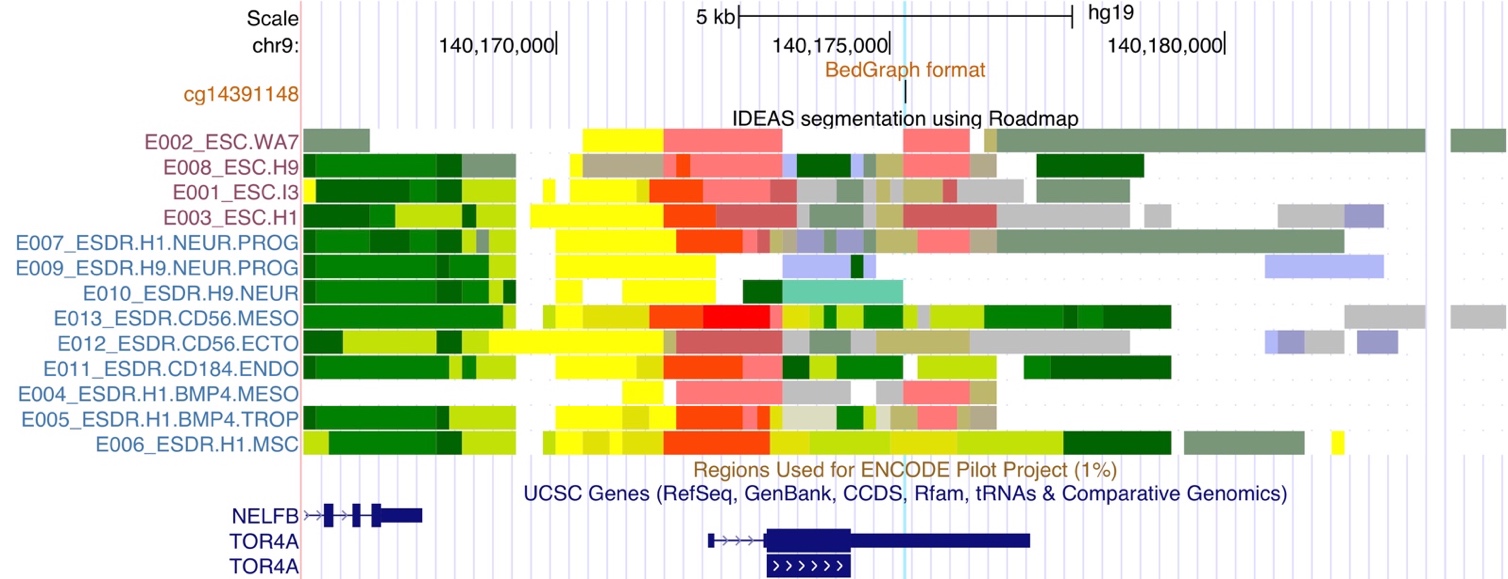

**Figure s4. Annotations for the genomic context of the five hub CpGs of the atypical POE module 3.** a: cg01331772; b: cg07274898; c: cg09639152; d: cg11464189; e: cg14391148. The color codes of IDEAS annotation using Roadmap data can be found in Figure 1a in Zhang et.al., 2017 (1).

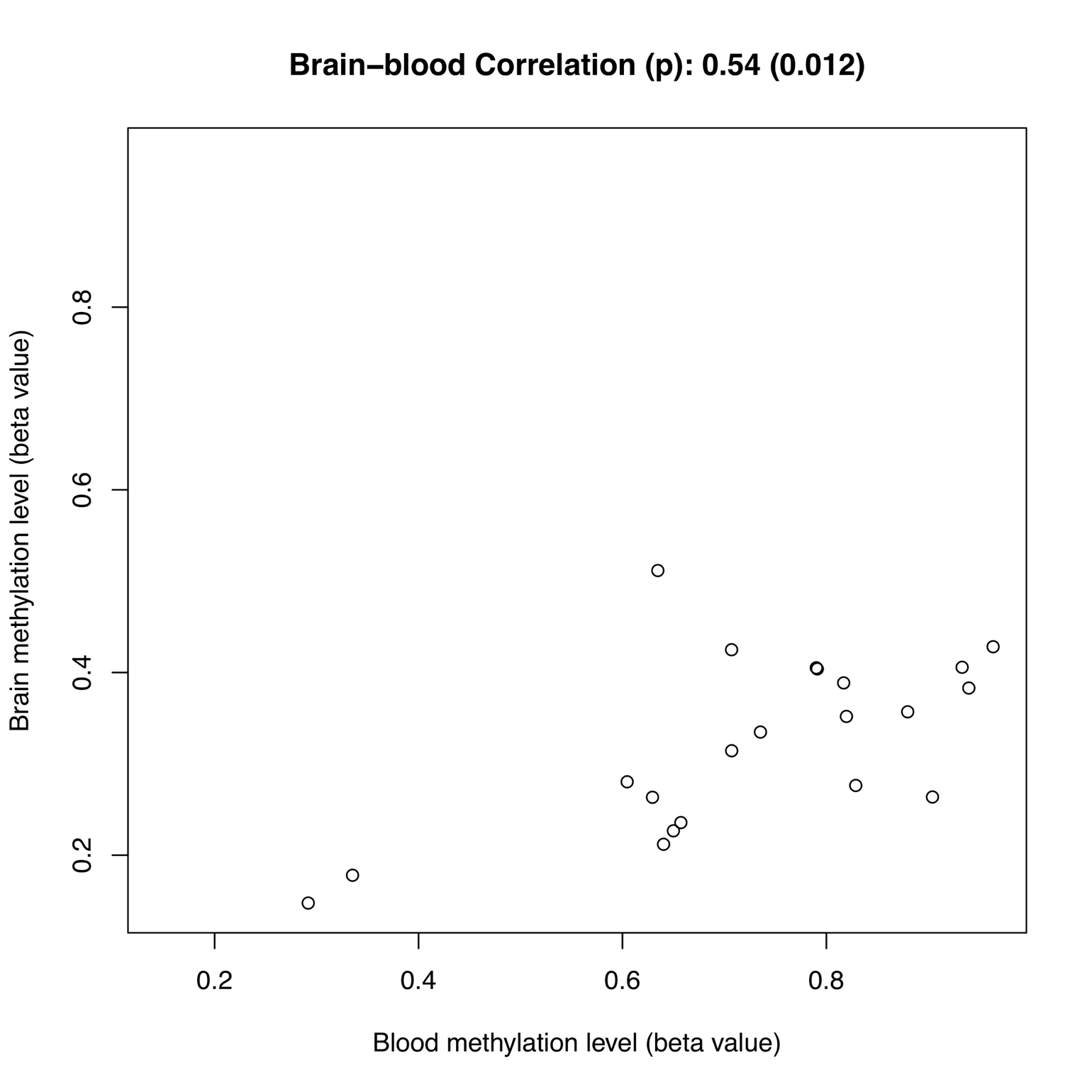

**Figure s5. The correlation between methylation levels of cg01331772 in blood and brain.** The results were extracted from IMAGE-CpG (2).

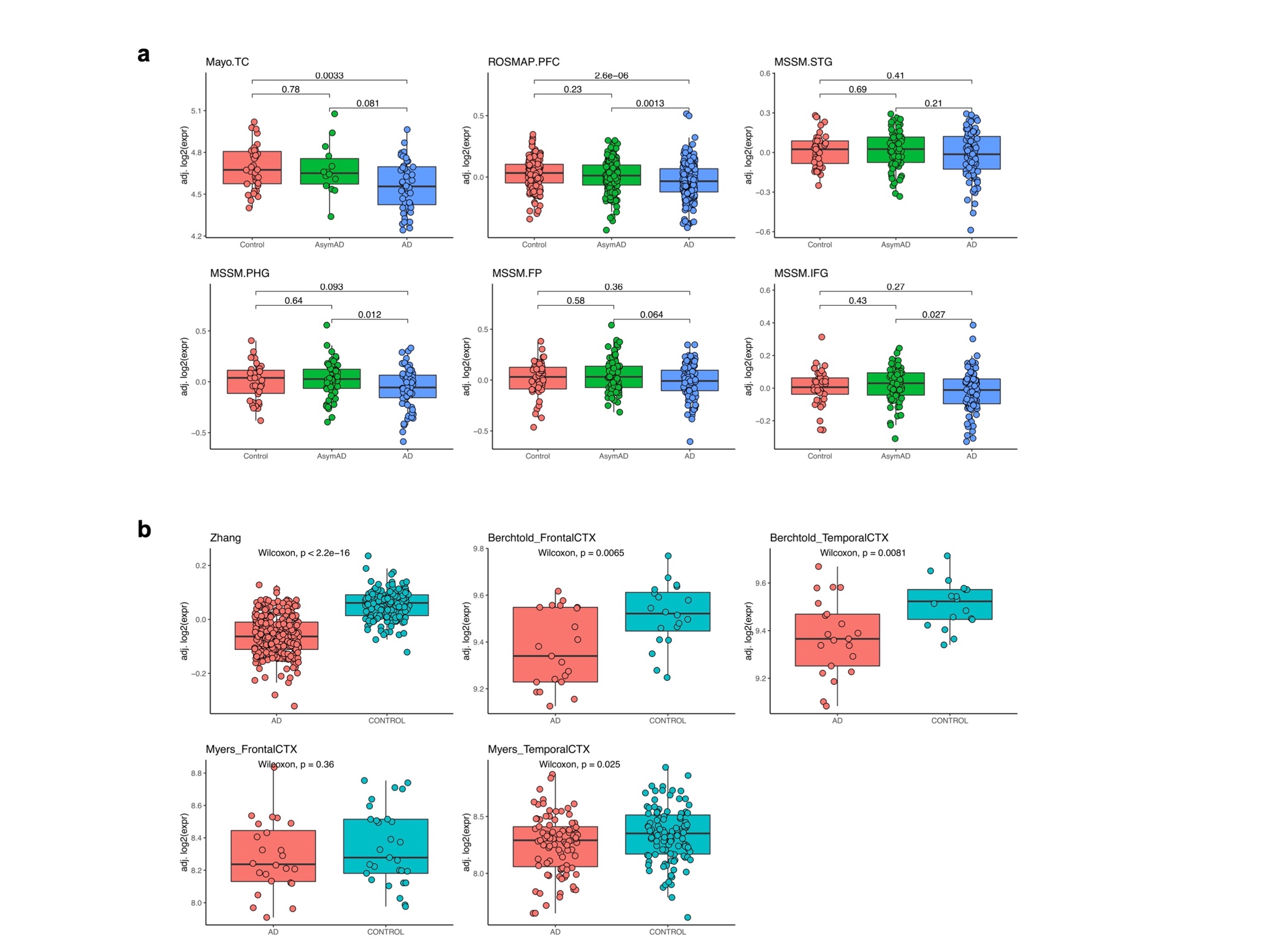

**Figure s6. Comparisons of *IMP4*'s mRNA expression in different brain tissues in control and Alzheimer's disease patients groups.** The results were extracted from http://swaruplab.bio.uci.edu:3838/bulkRNA/(3). a. Results from the consensus datasets. Mayo.TC: Mayo Clinic Brain Bank (Mayo) temporal cortex (TC); ROSMAP.PFC: Religious Orders Study and Memory and Aging Project (ROSMAP) prefrontal cortex (PFC); MSSM.STG/PHG/FP/IFG: Mount Sinai School of Medicine (MSSM) para-hippocampal gyrus (PHG), inferior frontal gyrus (IFG), superior temporal gyrus (STG) and frontal pole (FP). b. Results from the validation datasets. Zhang: prefrontal cortex in Zhang et al.(GSE44770); Berchtold_FrontalCTX/TemporalCTX: Frontal cortex and temporal cortex in PMID:23273601; Myers_FrontalCTX/TemporalCTX: Frontal cortex and temporal cortex in PMID:19361613
